# A mitochondrial SCF-FBXL4 ubiquitin E3 ligase complex restrains excessive mitophagy to prevent mitochondrial disease

**DOI:** 10.1101/2022.11.11.516094

**Authors:** Yu Cao, Jing Zheng, Huayun Wan, Yuqiu Sun, Song Fu, Shanshan Liu, Baiyu He, Gaihong Cai, Yang Cao, Huanwei Huang, Qi Li, Yan Ma, She Chen, Fengchao Wang, Hui Jiang

## Abstract

Mitophagy is a fundamental quality control mechanism of mitochondria. Its regulatory mechanisms and pathological implications remain poorly understood. Here via a mitochondria-targeted screen, we found that knockout (KO) of FBXL4, a mitochondrial disease gene, hyperactivates mitophagy at basal conditions. Subsequent counter screen revealed that FBXL4-KO hyperactivates mitophagy via two mitophagy receptors BNIP3 and NIX. We determined that FBXL4 functions as an integral outer-membrane protein that forms an SCF-FBXL4 ubiquitin E3 ligase complex. SCF-FBXL4 ubiquitinates BNIP3 and NIX to target them for degradation. Pathogenic FBXL4 mutations disrupt SCF-FBXL4 assembly and impair substrate degradation. *Fbxl4^−/−^* mice exhibit elevated BNIP3 and NIX proteins, hyperactive mitophagy, and perinatal lethality. Importantly, knockout of either *Bnip3* or *Nix* rescues metabolic derangements and viability of the *Fbxl4^−/−^* mice. Together, beyond identifying SCF-FBXL4 as a novel mitochondrial ubiquitin E3 ligase restraining basal mitophagy, our results reveal hyperactivated mitophagy as a cause of mitochondrial disease and suggest therapeutic strategies.

## INTRODUCTION

Mitophagy is an essential mitochondrial quality control mechanism that removes dysfunctional or excessive mitochondria by autophagy. Mitophagy defects associate with neurodegeneration, aging, cancer and other pathophysiological conditions (Moehlman & Youle, 2020, Onishi, Yamano et al., 2021, Youle & Narendra, 2011). Mammals have evolved multiple mitophagy pathways to remove mitochondria under various physiological and pathological conditions (Onishi et al., 2021, Youle & Narendra, 2011). For example, the Parkinson’s disease-related PINK1-Parkin pathway removes depolarized mitochondria and mitochondria with proteostatic stress (Narendra, Tanaka et al., 2008, Narendra, Jin et al., 2010, Pickrell & Youle, 2015). The mitophagy receptors BNIP3 and FUNDC1 mediate hypoxia-induced mitophagy (Liu, Feng et al., 2012, Zhang, Bosch-Marce et al., 2008). The mitophagy receptor NIX mediates mitochondria elimination during erythrocyte maturation (Sandoval, Thiagarajan et al., 2008, Schweers, Zhang et al., 2007). Moreover, characterizations of the mtKeima and mitoQC reporter mice reveal that mitophagy is widespread across tissues/organs at basal conditions (McWilliams, Prescott et al., 2016, McWilliams, Prescott et al., 2018, Sun, Yun et al., 2015). However, the regulatory mechanisms of basal mitophagy and their pathological implications remain poorly characterized.

The loss-of-function mutations of FBXL4 cause severe mitochondrial diseases characterized by mtDNA depletion, respiratory deficiency, lactic acidosis, encephalomyopathy and multisystem degeneration (Bonnen, Yarham et al., 2013, Gai, Ghezzi et al., 2013). FBXL4-KO mice exhibit excessive mitophagy, reduction of mitochondrial mass, and dominant perinatal lethality (Alsina, Lytovchenko et al., 2020). Despite the striking phenotype, the pathogenic mechanisms of FBXL4 mutations remain mysterious, and the mechanism of mitophagy activation and its pathological contribution in FBXL4-related diseases are unclear.

In this study, via a mitochondria-targeted CRISPR knockout screen for regulators of basal mitophagy, we identify that FBXL4-KO hyperactivates mitophagy in cells and *in vivo*. Subsequent genetic and biochemical characterizations reveal FBXL4 as a mitochondrial ubiquitin E3 ligase that restrains mitophagy via degrading mitophagy receptors BNIP3 and NIX. Knockout of either *Bnip3* or *Nix* represses hyperactive mitophagy, corrects metabolic derangements, and rescues viability of the *Fbxl4^−/−^* mice.

## RESULTS

### A mitochondria-targeted CRISPR knockout screen for mitophagy regulators

To monitor cellular mitophagy level, we exploited the mitoQC reporter, which expresses a GFP-mCherry tandem protein targeted to mitochondrial outer-membrane (Allen, Toth et al., 2013). Under normal condition, both GFP and mCherry are functional; when mitochondria are engulfed by lysosome, GFP is quenched by acidic pH but mCherry remains intact (**Figure 1A**). We established a HeLa tetracycline-inducible mitoQC (TO-mitoQC) reporter cell line, transformed the reporter line with Cas9 and a custom sgRNA library targeting the MitoCarta2.0 inventory of nuclear-encoded mitochondrial genes (**He et al., Cell Reports, accepted**), and induced mitoQC expression by doxycycline treatment. Mitophagy level was analyzed by FACS-based measurement of the mCherry/GFP fluorescence intensity ratio. Cells with enhanced (top 25% with high ratio) and reduced (bottom 25% with low ratio) mitophagy were sorted and sequenced (**Figure 1B**). Genes were ranked by calculating enrichment score (high-25%/low-25%) (**Figure 1C** and **Table S1**). We focused on four top hits with enhanced mitophagy (PPTC7, MFN2, FBXL4 and TMEM11) and one hit with reduced mitophagy (MARCH5) (highlighted in red, **Figure 1C**). Retesting with the mitoQC reporter and with two independent sgRNAs per gene verified the screen result (**Figure S1A**).

**Figure 1.**
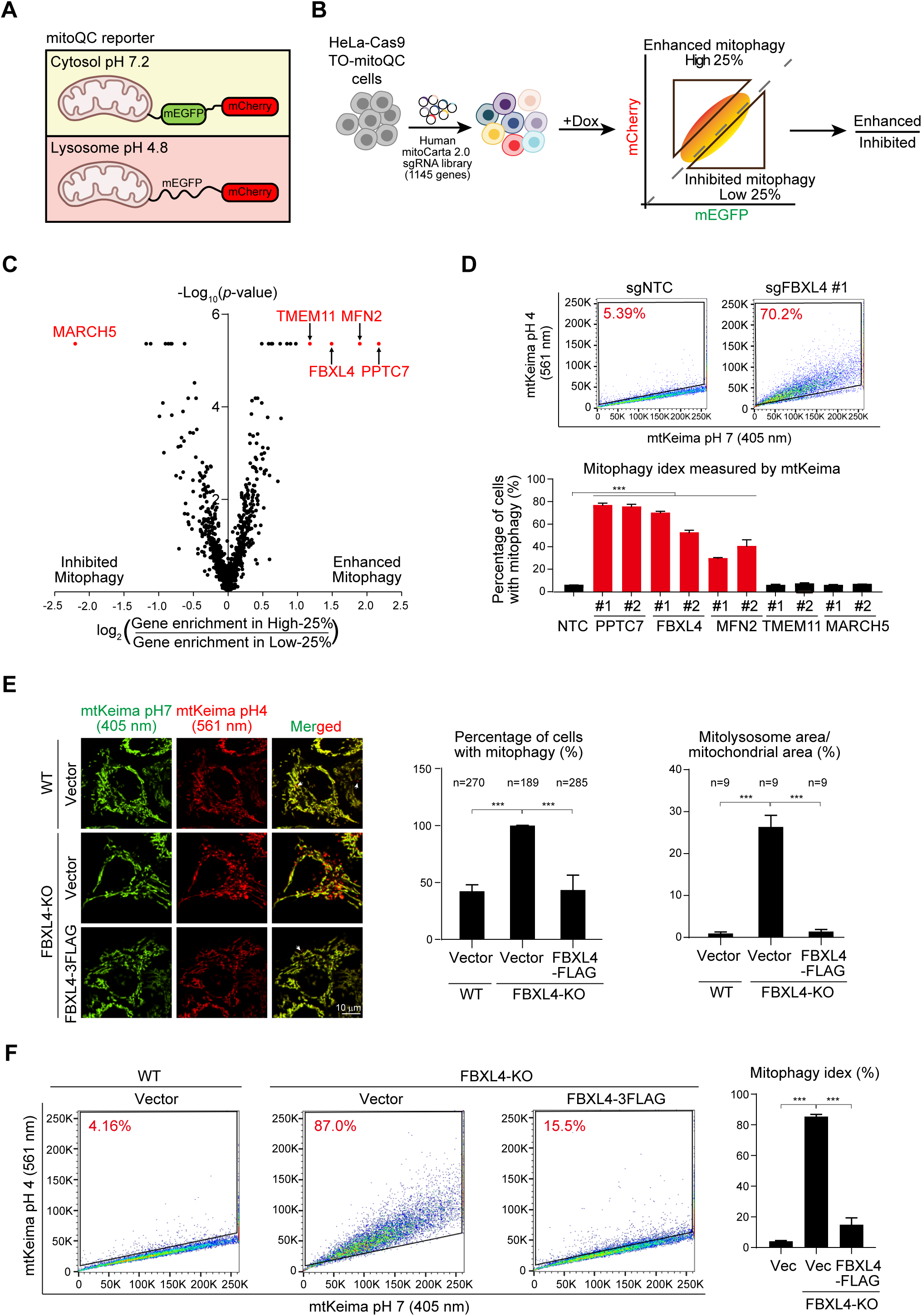
Mitochondria-targeted CRISPR-Cas9 screen for mitophagy regulators. (A) Schematic of the mitoQC reporter. (B) Schematic of the screen process of mitophagy regulators. (C) Volcano plot of the screen result. The top hits are highlighted in red. (D) Verification of screen hits by mtKeima-based FACS analysis of mitophagy. Upper: representative FACS analysis of mitophagy in HeLa cells expressing sgNTC and sgFBXL4. The percentage of cells with high mitophagy level is indicated in red. Bottom: quantitative analysis of mitophagy levels in HeLa cells expressing the indicated sgRNAs. Two independent sgRNAs were used for each gene. NTC: non-targeting control. (E) Imaging analysis of mitophagy levels in the indicated HeLa cells. Left: representative images, white arrows point to mitolysosomes (FBXL4-KO cells were not labeled because of the large numbers of mitolysosomes); middle: percentage of cells with mitophagy (mitolysosome-positive); right: quantification of mitolysosome/mitochondria area. n: number of cells analyzed (middle); number of imaging areas (20-30 cells/area) analyzed (right). (F) FACS analysis of mitophagy levels in the indicated HeLa cells. Left: representative FACS results; right: quantitative analysis of mitophagy levels. Data are mean + SD from three biological replicates (D, F). Statistics: two-tailed unpaired Student’s t-test (D-F); \*\*\**P* < 0.001.

We next analyzed mitophagy with the mtKeima reporter, a pH-sensitive protein targeted to mitochondrial matrix (Katayama, Kogure et al., 2011, Sun et al., 2015). Keima has an emission spectrum that peaks at 620 nm and a bimodal excitation at 440 nm (at pH 7.2) and at 586 nm (at pH 4.8) (**Figure S1B**). Mitophagy level was determined by FACS-based measurement of emissions from 561 nm excitation (acidic pH) and 405 nm excitation (neutral pH) (**Figure 1D**). Knockout of PPTC7, MFN2 and FBXL4 induced mitophagy but knockout of TMEM11 and MARCH5 did not (**Figure 1D**). The mitophagy phenotypes of TMEM11-KO and MARCH5-KO thus associate with the mitoQC reporter. This is likely because mitoQC induces higher basal mitophagy level than mtKeima (Liu, Sliter et al., 2021c). We thus used the mtKeima reporter in following studies.

### FBXL4 deficiency hyperactivates mitophagy

We focused on the hit FBXL4 because of its disease association and unknown pathological mechanism. We generated FBXL4-KO HeLa cells and confirmed its mitophagy phenotype. Fluorescence imaging with mtKeima showed that FBXL4-KO potently increased mitophagy (red-only mitolysosomes, **Figure 1E**). We confirmed the identity of mitolysosomes by showing their colocalization with the lysosome marker LAMP1-YFP (**Figure S1C**). Quantitative analysis showed that all the FBXL4-KO cells are mitophagy-positive whereas only ~45% wildtype (WT) cells are mitophagy-positive (**Figure 1E**). Mitolysosomes occupy ~25% of total mitochondria area in FBXL4-KO cells, in contrast to 0.9% in WT cells (**Figure 1E**). FACS analysis with mtKeima showed that the mitophagy level is <5% in WT cells but is increased to >80% in FBXL4-KO cells (**Figure 1F**). All these phenotypes of FBXL4-KO cells were rescued by re-expressing FBXL4-FLAG (**Figures 1E** and **1F**). Moreover, mitophagy in FBXL4-KO cells was blocked by knocking down FIP200 and Beclin1, two key autophagy genes (Mizushima & Komatsu, 2011) (**Figures S1D** and **S1E**).

We also generated another mitophagy reporter by anchoring mCherry to mitochondrial outer-membrane (**Figure S1F**). Lysosomal delivery of mitochondria causes reporter cleavage, releasing a free mCherry protein, such strategy was used in ER-phagy studies (Liang, Lingeman et al., 2018). Immunoblot analysis showed that ~15% of the reporter is cleaved in FBXL4-KO cells in contrast to 3.2% cleavage in WT cells and in FBXL4-KO cells rescued with FBXL4-FLAG (**Figures S1G**). Taken together, FBXL4-KO hyperactivates mitophagy in HeLa cells.

### FBXL4 deficiency post-transcriptionally upregulates BNIP3 and NIX to hyperactivate mitophagy

To reveal the molecular mechanism underlying hyperactive mitophagy in FBXL4-KO cells, we performed a counter screen with the MitoCarta2.0 sgRNA library, using mtKeima as the reporter. Our screen results showed that mitophagy induced by FBXL4-KO was inhibited by the knockout of two mitophagy receptors BNIP3 and NIX (**Figure 2A** and **Table S2**). Immunoblot analysis showed that FBXL4-KO greatly increased the protein levels of BNIP3 and NIX (**Figure 2B**). Interestingly, the mRNA levels of BNIP3 and NIX were reduced by 50% in FBXL4-KO cells (**Figure 2C**). The reduction of BNIP3 and NIX mRNAs in FBXL4-KO cells is likely an adaptive response to counteract hyperactivated mitophagy. This idea is supported by the observation that FBXL4-KO also downregulated another mitophagy receptor FUNDC1 (Liu et al., 2012) (**Figure 2B**). All the aberrant regulations of mitophagy receptors in FBXL4-KO cells were rescued by FBXL4-FLAG re-expression (**Figures 2B** and **2C**).

**Figure 2.**
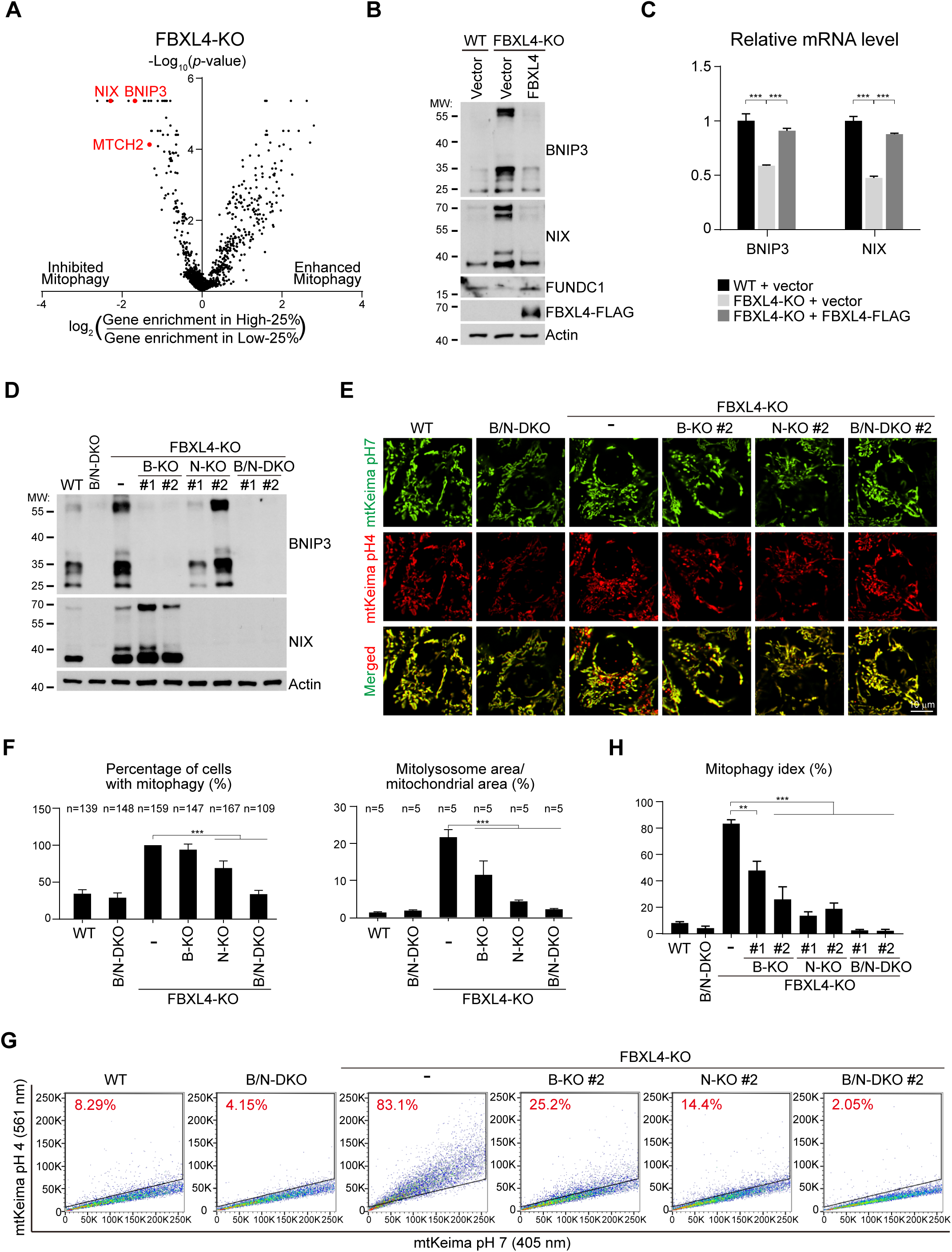
FBXL4 deficiency post-transcriptionally upregulates BNIP3 and NIX to hyperactivate mitophagy. (A) Volcano plot of the screen result of mitophagy regulators in FBXL4-KO HeLa cells. Mitochondrial outer-membrane protein MTCH2 and mitophagy receptors BNIP3 and NIX are highlighted in red. (B) Immunoblot analysis of the indicated HeLa cells. WT and FBXL4-KO HeLa cells were rescued with vector or FBXL4-FLAG. (C) qPCR analysis of BNIP3 and NIX in the indicated HeLa cells. (D) Immunoblot analysis of the indicated HeLa cells. B: BNIP3; N: NIX. Two clones were shown for each double and triple knockout cells. (E and F) Representative live cell imaging (E) and quantitative analysis (F) of mitophagy levels in the indicated HeLa cells. n: number of cells analyzed (left); number of imaging areas analyzed (right). (G) Representative FACS analysis of mitophagy levels in the indicated HeLa cells. (H) FACS-based quantitative analysis of mitophagy levels in the indicated HeLa cells. Data are mean + SD from three biological replicates (C, H). Statistics: two-tailed unpaired Student’s t-test (C, F and H); \*\**P* < 0.01; \*\*\**P* < 0.001.

We next knocked out BNIP3 and NIX individually or combinatorically in WT and FBXL4-KO HeLa cells (**Figure 2D**). Fluorescence imaging and FACS analysis with the mtKeima reporter showed that individual knockout of BNIP3 and NIX partially inhibited mitophagy, and double knockout of BNIP3 and NIX completely blocked mitophagy in FBXL4-KO cells (**Figures 2E-2H**). Therefore, FBXL4-KO hyperactivates mitophagy via BNIP3 and NIX.

MTCH2 encodes a mitochondrial outer-membrane protein regulating apoptosis and metabolism (Gross, 2016) and is recently identified as a mitochondrial outer-membrane insertase for α-helical proteins (Guna, Stevens et al., 2022). MTCH2-KO also inhibited mitophagy in FBXL4-KO cells (**Figures 2A**, **S2A** and **S2B**). Immunoblot showed that MTCH2-KO reduced the protein levels of BNIP3 and NIX in both WT and FBXL4-KO cells (**Figure S2C**). These results support BNIP3 and NIX as the central node to regulate excessive mitophagy in FBXL4-KO cells.

### FBXL4 is an integral mitochondrial outer-membrane protein

To understand how FBXL4 regulates BNIP3 and NIX, we firstly determined the submitochondrial localization of FBXL4. We purified mitochondria from HeLa cells stably expressing FBXL4-FLAG and performed Protease K digestion with different mitochondrial preparations. With intact mitochondria, Protease K only digests outer-membrane proteins exposed to the cytosol (Tom70). Swelling with hypotonic buffer disrupts mitochondrial outer-membrane but leaves inner membrane intact, resulting in the cleavage of inter-membrane space proteins (Smac) and inner membrane proteins exposed to the inter-membrane space (Mitofilin). After lysis with Triton X-100, both mitochondrial membranes are lysed, resulting in the cleavage of all the mitochondrial proteins, including matrix proteins (Hsp60). FBXL4-FLAG was cleaved by Protease K digestion in intact mitochondria preparation (lane 2, **Figure 3A**). We repeated digestion of intact mitochondria with trypsin and found that FBXL4-FLAG was also cleaved (lane 7, **Figure 3A**). These results suggest FBXL4 as an outer-membrane protein.

**Figure 3.**
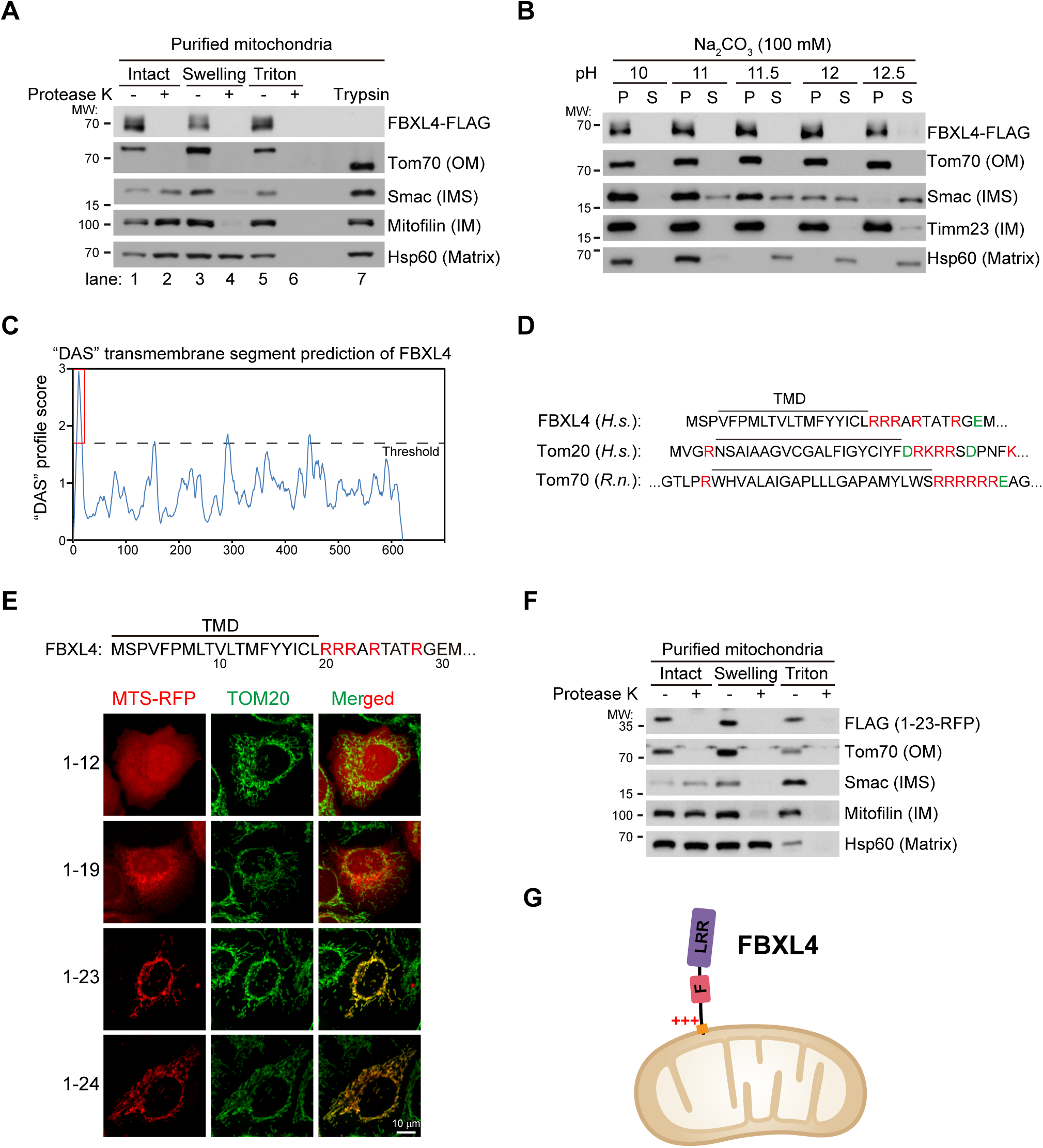
FBXL4 is an integral mitochondrial outer-membrane protein. (A) Determination of the submitochondrial localization of FBXL4 by Protease K and trypsin digestion. Mitochondria from HeLa FBXL4-FLAG stable line were purified and stored as intact mitochondria or treated with hypotonic swelling buffer or lysed with Triton X-100 buffer. Different mitochondrial preparations were then digested with Protease K or trypsin. (B) Analysis of the association of FBXL4 with membrane by alkaline carbonate extraction. Purified mitochondria from HeLa FBXL4-FLAG stable line were treated with alkaline carbonate buffer at the indicated pH and then centrifuged to collect the supernatant and the pellet fractions. S: supernatant; P: pellet. (C) “DAS” prediction of the transmembrane domain (TMD) of FBXL4. The red box highlights the putative TMD at FBXL4 N-terminus. (D) Analysis of the TMD and the flanking sequences of FBXL4, Tom20 and Tom70. Positively-charged residues are highlighted in red; negatively-charged residues are highlighted in green. *H.s.*: Homosapien; *R.n.*: Rattus norvegicus. (E) Identification of the mitochondrial targeting sequence of FBXL4. Indicated FBXL4 N-terminal sequences were fused to RFP and expressed in HeLa cells to analyze their subcellular localization. (F) Determination of the submitochondrial localization of FBXL4(1-23)-RFP by Protease K digestion. Experiments were performed similarly as (A).

To determine if FBXL4 is an integral membrane protein or peripherally associates with mitochondria, we purified mitochondria and performed alkaline extraction (Fujiki, Hubbard et al., 1982). Soluble and peripheral proteins will be extracted to the soluble fraction by alkaline pH, whereas membrane proteins remain integral to the membrane and appear in the pellet fraction. At increasing pH from 10 to 12.5, FBXL4-FLAG remained in the pellet (mitochondria) like the integral membrane proteins Tom70 and Tim23 (**Figure 3B**). In contrast, soluble proteins Smac and Hsp60 were extracted to the supernatant fraction (**Figure 3B**). Therefore, FBXL4 is an integral mitochondrial outer-membrane protein.

### Determination of the transmembrane domain, targeting sequence and topology of FBXL4

Analysis of FBXL4 protein sequence with the “DAS” transmembrane prediction server (https://tmdas.bioinfo.se/) predicted that FBXL4 contains a putative transmembrane domain (TMD) at its N-terminus (highlighted in red box, **Figure 3C**). We noticed that the putative TMD is followed by a stretch of positively-charged arginine residues, a typical feature of N-terminally anchored mitochondrial outer-membrane proteins, such as Tom20 and Tom70 (Rapaport, 2003) (**Figure 3D**).

To determine if the N-terminal sequence of FBXL4 truly forms a mitochondrial outer-membrane targeting sequence, we fused variable truncations of FBXL4 N-terminal sequence (amino acids 1-24) to RFP and analyzed the subcellular localization of the fusion proteins. Immunofluorescence imaging showed that amino acids 1-12 targeted RFP to the cytosol, 1-19 (TMD) targeted RFP to non-mitochondrial vesicular structures, whereas 1-23 and 1-24 (TMD + arginine stretch) targeted RFP to mitochondria (**Figure 3E**). Protease K digestion experiment showed that FBXL4(1-23)-RFP localized to mitochondrial outer-membrane (**Figure 3F**). Therefore, FBXL4 is an N-anchored mitochondrial outer-membrane protein targeted by amino acids 1-23 (**Figure 3G**).

### FBXL4 forms an SCF-FBXL4 ubiquitin E3 ligase complex and associates with the UBXD8-VCP complex

FBXL4 contains an F-box domain and a C-terminal leucine rich repeats (LRR) (**Figure 4A**). F-box is a protein domain that interacts with the Skp1 adaptor to form an Skp1-Cullin1-F-box (SCF) ubiquitin E3 ligase complex (Bai, Sen et al., 1996, Petroski & Deshaies, 2005). We aligned the F-box sequence of FBXL4 with that of multiple canonical F-box proteins (Jin, Cardozo et al., 2004) and found that the four residues critical for Skp1 binding (Bai et al., 1996) are perfectly conserved in FBXL4 (**Figure 4A**).

**Figure 4.**
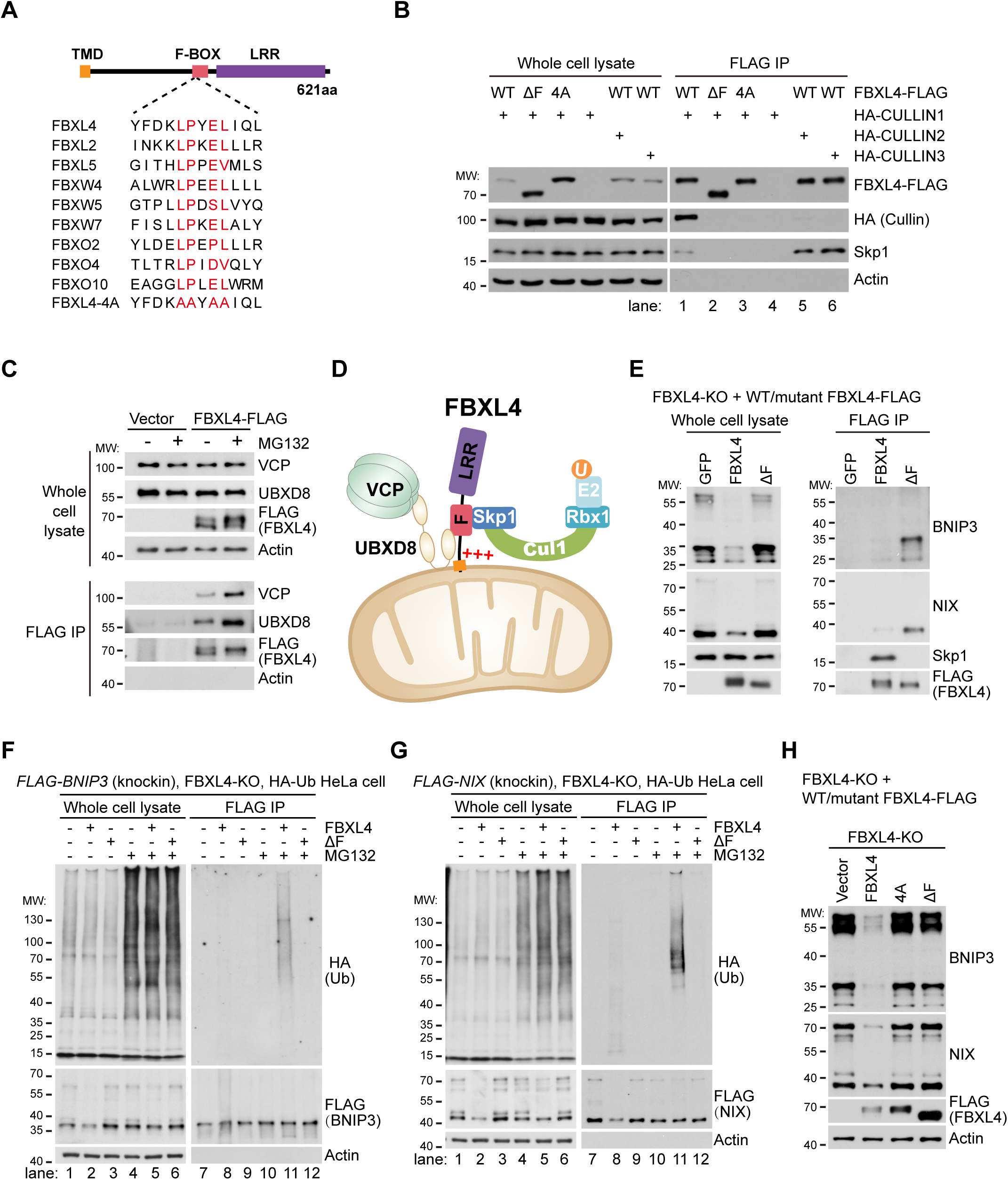
FBXL4 forms an SCF-FBXL4 ubiquitin E3 ligase complex to ubiquitinate and degrade BNIP3 and NIX. (A) Alignment of F-box sequences. Critical residues for Skp1 binding are highlighted in red. (B) Immunoprecipitation analysis of the FBXL4-Skp1-Cullin1 (SCF-FBXL4) complex. HeLa cells were infected with lentiviruses expressing the indicated genes. ΔF: the F-box deletion mutant of FBXL4-FLAG; 4A: FBXL4-4A mutant as shown in (A). (C) Immunoprecipitation analysis the FBXL4-UBXD8-VCP complex. HeLa cells expressing vector or FBXL4-FLAG were treated with DMSO or MG132 (20 μM) for 8 hours. (D) Schematic of the SCF-FBXL4-UBXD8-VCP complex at mitochondrial outer-membrane. (E) Immunoprecipitation analysis of the association of FBXL4 with BNIP3 and NIX. FBXL4-KO HeLa cells were rescued with GFP, FBXL4-FLAG or FBXL4(ΔF)-FLAG. (F) Immunoprecipitation analysis of the ubiquitination of BNIP3. FLAG-BNIP3 (knockin), FBXL4-KO, HA-ubiquitin (Ub) HeLa cells were rescued with WT or ΔF FBXL4, and treated with DMSO or MG132 (20 μM) for 8 hours. (G) Immunoprecipitation analysis of the ubiquitination of NIX. FLAG-NIX (knockin), FBXL4-KO, HA-Ub HeLa cells were treated the same as (F). (H) Immunoblot analysis of the indicated HeLa cells. FBXL4-KO HeLa cells were rescued with vector, FBXL4-FLAG, FBXL4(4A)-FLAG or FBXL4(ΔF)-FLAG.

To determine if FBXL4 forms an SCF ubiquitin E3 ligase complex, we expressed WT or F-box mutant FBXL4-FLAG, and HA-tagged Cullin1-3 in HeLa cells. Anti-FLAG immunoprecipitation (Platt, Chen et al.) of FBXL4-FLAG pulled down endogenous Skp1 and HA-Cullin1 but not HA-Cullin2 or HA-Cullin3 (lane 1 vs. lanes 5 & 6, **Figure 4B**). Deleting the F-box (ΔF) or mutating the four key residues of F-box to alanine (4A) completely abolished FBXL4 interaction with Skp1 and HA-Cullin1 (lane 1 vs. lanes 2 & 3, **Figure 4B**). We recently reported that mitochondrial UBXD8-VCP complex associates with mitochondrial ubiquitin E3 ligases and mediates the degradation of substrates, including BNIP3 (Zheng, Cao et al., 2022). Anti-FLAG IP of FBXL4-FLAG pulled down UBXD8 and VCP at normal and proteasome inhibition (MG132 treatment) conditions (**Figure 4C**). Taken together, FBXL4 forms an SCF-FBXL4 ubiquitin E3 ligase complex and associates with the UBXD8-VCP complex at mitochondrial outer-membrane (**Figure 4D**).

### The SCF-FBXL4 ubiquitin E3 ligase complex mediates the ubiquitination and degradation of BNIP3 and NIX

F-box protein is the substrate-recruiting adaptor in the SCF ubiquitin E3 ligase complex. We thus examined if FBXL4 associates with BNIP3 and NIX. We rescued FBXL4-KO HeLa cells with WT or ΔF FBXL4-FLAG. Anti-FLAG IP showed that WT FBXL4-FLAG pulled down Skp1 but very few BNIP3 or NIX (**Figure 4E**), likely because the two substrates are degraded. In contrast, FBXL4(ΔF)-FLAG clearly pulled down BNIP3 and NIX but not Skp1 (**Figure 4E**). Notably, protein samples from IP experiments lose the dimeric forms of BNIP3 and NIX on immunoblot (**Figure 4E**). This is not an artifact because we found that quick processing of fresh cell samples preserves the dimeric forms of BNIP3 and NIX as shown in **Figure 2B**. But preserving cells/tissues for extended time at −80 °C and prolonged cell lysate incubation at 4 °C during IP experiment cause the loss of dimeric forms, which is evident in **Figure 4E** and also in **Figures 4F**, **5D**, **6C**, **7C**, **S5A** and **S5B**.

**Figure 5.**
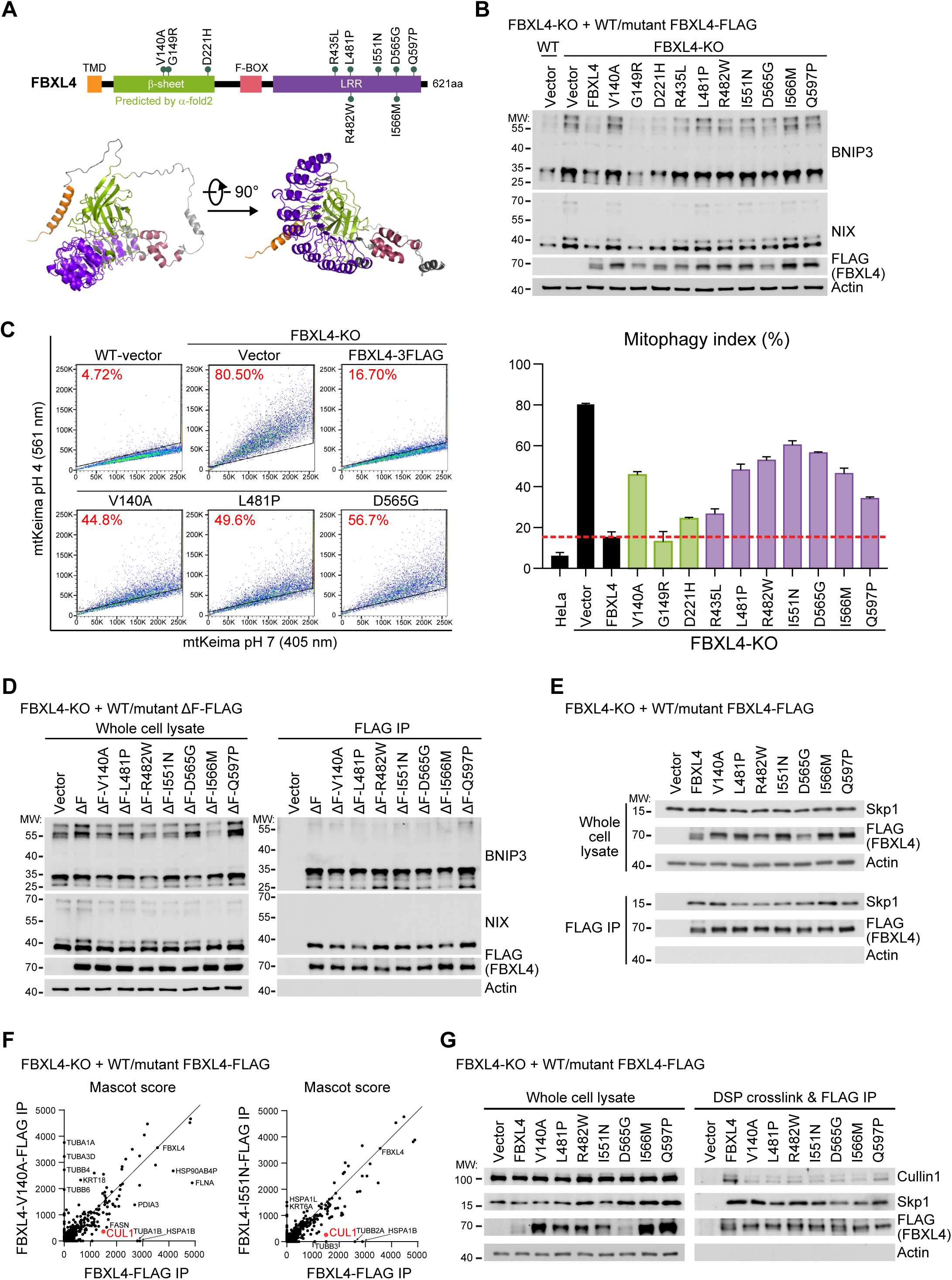
Pathogenic mutations of FBXL4 disrupt SCF-FBXL4 complex assembly and impair the degradation of BNIP3 and NIX to hyperactivate mitophagy. (A) Schematic of FBXL4 domain organization and predicted structure by AlphaFold2. (B) Immunoblot analysis of the indicated HeLa cells. WT and FBXL4-KO HeLa cells were rescued with vector, WT or mutant FBXL4-FLAG. Functionally-defective FBXL4 mutants are highlighted in blue. (C) FACS-based analysis of mitophagy levels in the indicated HeLa cells. The same HeLa cells in (B) were used. Left: representative FACS analysis; right: quantitative analysis of mitophagy levels. The dashed red line marks the mitophagy level in FBXL4-KO cells rescued with FBXL4-FLAG. Data are mean + SD from three biological repeats. (D) Immunoprecipitation analysis of FBXL4-substrate interaction. FBXL4-KO HeLa cells were rescued with vector, FBXL4(ΔF)-FLAG or mutant FBXL4(ΔF)-FLAG. (E) Immunoprecipitation analysis of FBXL4-Skp1 interaction. FBXL4-KO HeLa cells were rescued with vector, WT or mutant FBXL4-FLAG. (F) Analysis of the mass spectrometry results of immunoprecipitation with WT or mutant FBXL4-FLAG. FBXL4-KO HeLa cells expressing WT, V140A or I551N FBXL4-FLAG were subject to DSP crosslinking and anti-FLAG immunoprecipitation. The immunoprecipitates were analyzed by mass spectrometry. The Mascot score of immunoprecipitated proteins were plotted. Because of space limitation, 6 common proteins with Mascot score between 5,000-20,000 were removed. Cullin1, which has decreased interaction with both FBXL4 mutants, is highlighted in red. (G) Immunoprecipitation analysis of the integrity of the SCF-FBXL4 complex. FBXL4-KO HeLa cells expressing vector, WT or mutant FBXL4-FLAG were crosslinked with DSP and subject to anti-FLAG immunoprecipitation.

To determine if SCF-FBXL4 ubiquitinates BNIP3 and NIX, we knocked in a 3×FLAG tag to the BNIP3 or NIX locus in FBXL4-KO HeLa cells. These knockin cell lines allow us to pull down BNIP3 and NIX expressed at endogenous level. We rescued the FLAG-BNIP3/NIX (knockin), FBXL4-KO, HA-ubiquitin lines with WT or ΔF FBXL4 and treated cells with DMSO or MG132. IP of FLAG-BNIP3 and FLAG-NIX showed that although both proteins accumulate in FBXL4-KO cells, they are not ubiquitinated under basal and MG132-treated conditions (lanes 7 & 10, **Figures 4F** and **4G**). MG132 treatment stabilized ubiquitinated forms of FLAG-BNIP3 and FLAG-NIX in cells expressing WT but not ΔF FBXL4 (lane 11 vs. lane 12, **Figures 4F** and **4G**). These results demonstrate that FBXL4 ubiquitinates BNIP3 and NIX.

The elevated protein levels of BNIP3 and NIX in FBXL4-KO cells was rescued by WT FBXL4 but not by its F-box mutants (**Figure 4H**), suggesting that other SCF subunits are essential for FBXL4 function. To examine the role of Cullin1, we inducibly overexpressed a dominant-negative form of Cullin1 (DN-Cullin1), which lacks the C-terminal Rbx1 binding region (Wu, Fuchs Serge et al., 2000). DN-Cullin1 overexpression did not affect the mRNA levels of BNIP3 and NIX but increased their protein levels in WT cells but not in FBXL4-KO cells (**Figures S3A** and **S3B**). Functionally, overexpression of DN-Cullin1 activated mitophagy in WT cells but not in BNIP3/NIX-DKO cells (**Figures S3C** and **S3D**), demonstrating that Cullin1 inactivation activates mitophagy via BNIP3 and NIX. Taken together, these results demonstrate that BNIP3 and NIX are substrates of the SCF-FBXL4 complex.

### Pathogenic mutations of FBXL4 disrupt the assembly of the SCF-FBXL4 complex and impair the degradation of BNIP3 and NIX to hyperactivate mitophagy

To examine if patient-derived mutations of FBXL4 affect mitophagy, we collected 10 mutations from literature (Barøy, Pedurupillay et al., 2016, El-Hattab, Dai et al., 2017, Gai et al., 2013, Huemer, Karall et al., 2015) (**Figure 5A**). Because three mutations (V140A, G149R and D221H) localize to a region without domain annotation, we predicted 3D structure of FBXL4 by AlphaFold2 (Jumper, Evans et al., 2021). Interestingly, AlphaFold2 predicted an unnamed domain rich in β-sheet. We thus temporarily named this domain as “β-sheet” (shown in green, **Figure 5A**). All the FBXL4 mutations localize either to the β-sheet (V140A, G149R and D221H) or the LRR domain (R435L, L481P, R482W, I551N, D565G, I566M and Q597P). We rescued FBXL4-KO cells with WT or mutant FBXL4-FLAG. All the patient-derived mutants, except G149R, failed to reduce BNIP3 and NIX levels (highlighted in blue, **Figure 5B**) and could not repress hyperactivated mitophagy in FBXL4-KO cells (**Figure 5C**). We then rescued the FLAG-BNIP3 (knockin), FBXL4-KO, HA-ubiquitin cells with WT or mutant FBXL4, treated cells with MG132 to block FLAG-BNIP3 degradation, and immunoprecipitated FLAG-BNIP3 to examine its ubiquitination status. FLAG-BNIP3 ubiquitination was apparently much less in cells expressing mutant FBXL4 as compared to cells expressing WT FBXL4 (**Figure S4**). These results demonstrate that FBXL4 pathogenic mutants are defective in substrate ubiquitination and degradation.

To understand how these pathogenic mutations affect SCF-FBXL4 function, we firstly examined substrate binding. We introduced these mutations to FBXL4(ΔF)-FLAG, which has stable interaction with substrates (**Figure 4E**), and rescued FBXL4-KO cells with WT or mutant FBXL4(ΔF)-FLAG (**Figure 5D**). IP of WT and mutant FBXL4(ΔF)-FLAG pulled down similar amounts of BNIP3 and NIX (**Figure 5D**), indicating FBXL4 pathogenic mutations do not interfere with substrate binding.

We next examined if the mutations affect FBXL4 interaction with Skp1. We rescued FBXL4-KO cells with WT or mutant FBXL4-FLAG (**Figure 5E**). IP of WT and mutant FBXL4-FLAG pulled down similar amounts of Skp1 (**Figure 5E**).

Given that SCF-FBXL4 is a membrane protein complex that may have unidentified binding partners, we performed chemical crosslinking with the reversible crosslinker DSP to stabilize the protein complexes in cells (Kim, Sarbassov et al., 2002) and then performed anti-FLAG IP of WT, V104A and I551N FBXL4-FLAG. Mass spectrometry analysis of immunoprecipitates identified Cullin1 as the only common protein that had decreased interaction with both V104A and I551N FBXL4-FLAG (**Figure 5F** and **Table S3**), indicating the two pathogenic mutations impair the assembly of the SCF-FBXL4 complex. We thus analyzed other FBXL4 mutants by DSP crosslinking and immunoprecipitation. We were surprised to find that all these mutants pulled down similar levels of Skp1 but less Cullin1 than WT FBXL4-FLAG (**Figure 5G**). Together, these results suggest that FBXL4 pathogenic mutations disrupt the assembly of the SCF-FBXL4 complex to impair the degradation of BNIP3 and NIX.

### *Fbxl4* deficiency increases the protein levels of BNIP3 and NIX and hyperactivates mitophagy *in vivo*

We generated and characterized the *Fbxl4*-null mice to investigate if FBXL4 regulates BNIP3 and NIX, as well as mitophagy *in vivo*. *Fbxl4^+/-^* mice were normal, but ~80% of the *Fbxl4^−/−^* mice died within 3 days after birth (**Figure 6A**). We thus collected tissue samples from P0 mice. Immunoblot analysis detected the accumulation of BNIP3 and NIX in *Fbxl4^−/−^* liver, heart, kidney and muscle (**Figure 6B**). Only BNIP3 accumulated in *Fbxl4^−/−^* lung, whereas neither BNIP3 nor NIX accumulated in *Fbxl4^−/−^* brain (**Figure 6B**). Consistent with the accumulation of BNIP3 and NIX, we observed decreased levels of mitochondrial proteins in *Fbxl4^−/−^* liver, heart, kidney, muscle and lung (**Figure 6B**), indicating the reduction of mitochondrial mass.

**Figure 6.**
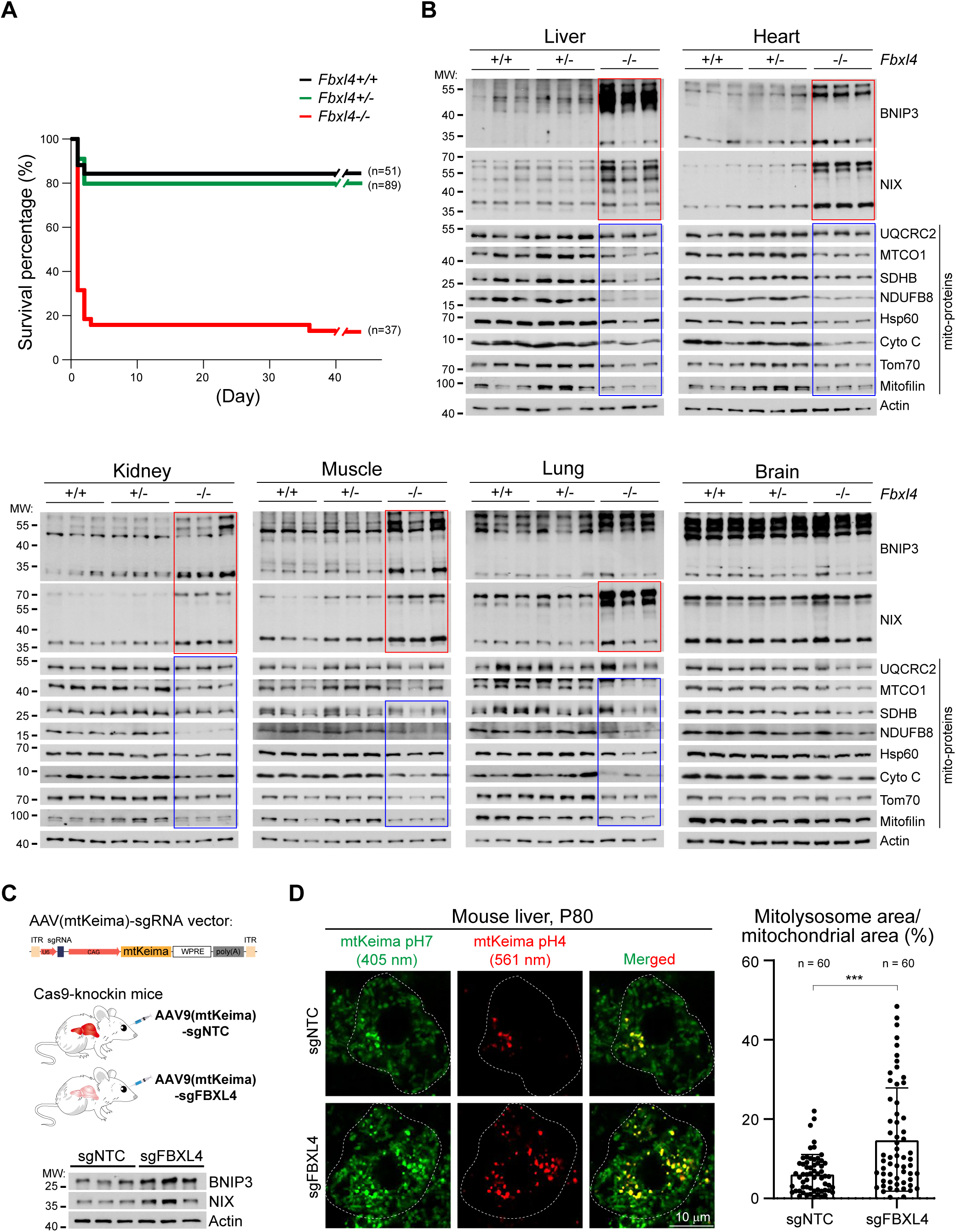
*Fbxl4* deficiency increases the protein levels of BNIP3 and NIX and hyperactivates mitophagy *in vivo* in mice. (A) Viability of the indicated mice. n: number of mice analyzed. (B) Immunoblot analysis of organs from the indicated mice at P0 of age. Elevated BNIP3 and NIX levels are highlighted by red boxes; reduced mitochondrial proteins are highlighted by blue boxes. Three mice for each genotype were analyzed. (C) Schematic of AAV-based gene knockout and mitophagy examination. Upper: schematic of the bicistronic AAV(mtKeima)-sgRNA vector that simultaneously expresses mtKeima and sgRNA; Middle: schematic of delivering AAV9(mtKeima)-sgNTC or AAV9(mtKeima)-sgFBXL4 to the Cas9-knockin mice to generate the control mice and hepatic *Fbxl4*-KO mice; below: immunoblot analysis of the indicated mouse liver. Three mice for each genotype were analyzed. (D) Live cell confocal imaging of mitophagy level in the indicated mouse liver. Left: representative images. The dashed line indicates cell boundary. Right: quantitative analysis of mitophagy levels in hepatocytes. n: number of cells analyzed. Data are mean ± SD. Statistics: two-tailed unpaired Student’s t-test; \*\*\**P* < 0.001.

Patient-derived fibroblasts exhibited enhanced mitophagy as measured by the mitoQC reporter (Alsina et al., 2020), but whether *Fbxl4*-KO hyperactivates mitophagy *in vivo* remains unclear. We did not choose to analyze the mitophagy level of the living *Fbxl4^−/−^* mice because these mice may have unknown adaptations. Instead, we exploited the CRISPR-Cas9 technology to knockout *Fbxl4* in adult mice (Liu, Fu et al., 2021b, Platt et al., 2014). We engineered an adeno-associated virus (AAV) vector that dually expresses the mtKeima reporter and sgRNA (AAV(mtKeima)-sgRNA) (**Figure 6C**). We delivered AAV9(mtKeima)-sgFBXL4 or AAV9(mtKeima)-sgNTC (NTC: non-targeting control) to the Cas9-knockin mice (**Figure 6C**). Due to the lack of FBXL4 antibody, we could not probe *Fbxl4* knockdown efficiency. But immunoblot analysis of mouse liver samples showed that AAV9(mtKeima)-sgFBXL4 elevated the protein levels of BNIP3 and NIX after 45 days of AAV delivery (**Figure 6C**), suggesting sgFBXL4 works. Notably, the tissue samples were cryopreserved for extended time and lost the dimeric forms of BNIP3 and NIX. We thus did not show the high molecular weight region in the immunoblots of BNIP3 and NIX to save space.

We next dissected fresh liver samples and imaged mtKeima by confocal microscopy. Mouse livers infected with AAV9(mtKeima)-sgFBXL4 exhibited significantly higher mitophagy levels than livers infected with AAV9(mtKeima)-sgNTC (**Figure 6D**). Collectively, these results demonstrate that *Fbxl4*-KO increases the protein levels of BNIP3 and NIX and hyperactivates mitophagy *in vivo*.

### Knockout of either *Bnip3* or *Nix* rescues mitochondrial content, metabolic derangements and viability of the *Fbxl4*-null mice

To determine if hyperactivated mitophagy causes perinatal lethality of the *Fbxl4^−/−^* mice, we knocked out *Bnip3* and *Nix*. Strikingly, ~60% of the *Fbxl4^−/−^ Bnip3^+/-^* and *Fbxl4^−/−^ Nix^+/-^* mice, and ~80% of the *Fbxl4^−/−^ Bnip3^−/−^* and *Fbxl4^−/−^ Nix^−/−^* mice survived to the adulthood (**Figure 7A**). The oldest *Fbxl4^−/−^ Bnip3^−/−^* mice were over 280 days of age at manuscript preparation. We also bred the *Bnip3^−/−^ Nix^−/−^* mice but these mice were unhealthy and infertile. We thus did not try breeding the *Fbxl4^−/−^ Bnip3^−/−^ Nix^−/−^* mice.

**Figure 7.**
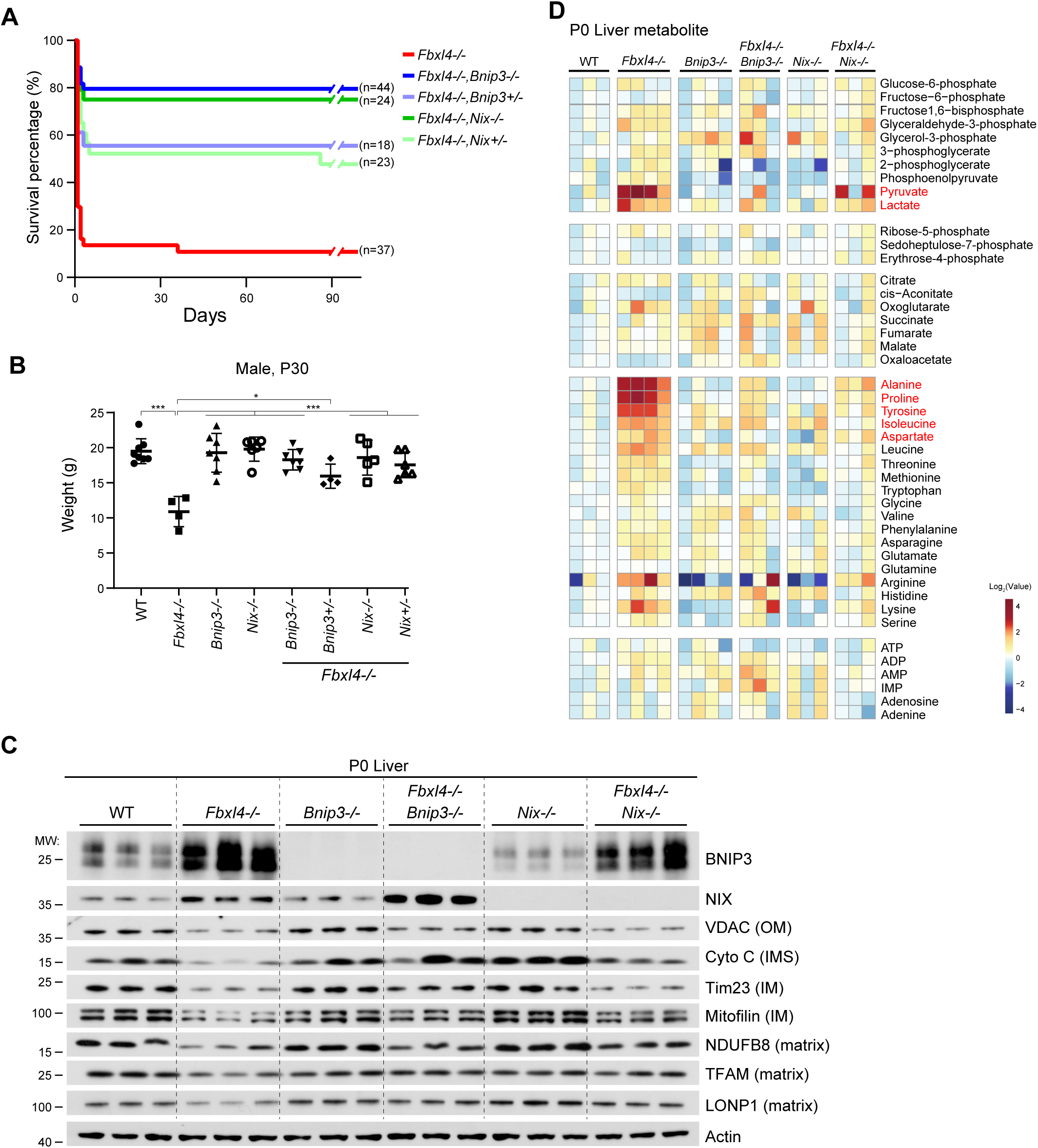
Knockout of either *Bnip3* or *Nix* rescues the *Fbxl4^−/−^* mice. (A) Viability of the indicated mice. n: number of mice analyzed. (B) Body weight of the indicated male mice at P30. Data are mean ± SD. Statistics: one-way ANOVA with the Tukey-Kramer test; *p < 0.05, ***p < 0.001. (C) Immunoblot analysis of liver samples from the indicated mice at P0 of age. Three mice for each genotype were analyzed. (D) Metabolite analysis of liver samples from the indicated mice at P0 of age. Three to five mice for each genotype were analyzed.

Most of the *Fbxl4^−/−^* mice died perinatally, and the <20% living *Fbxl4^−/−^* mice were much smaller than WT mice at P30 (**Figure 7B**), indicating developmental delay. Remarkably, the *Fbxl4^−/−^ Bnip3^+/-^*, *Fbxl4^−/−^ Nix^+/-^*, *Fbxl4^−/−^ Bnip3^−/−^* and *Fbxl4^−/−^ Nix^−/−^* mice had similar body weight as WT mice at P30 (**Figure 7B**). Therefore, reducing BNIP3 or NIX rescues not only perinatal lethality but also developmental delay of the *Fbxl4^−/−^* mice.

We next collected tissue samples from WT, *Fbxl4^−/−^*, *Bnip3^−/−^*, *Fbxl4^−/−^ Bnip3^−/−^*, *Nix^−/−^* and *Fbxl4^−/−^ Nix^−/−^* mice at P0. Immunoblot analysis of liver samples showed that *Bnip3*-KO is better than *Nix*-KO to rescue mitochondrial protein levels of the *Fbxl4^−/−^* mice, especially in the cases of VDAC, Cytochrome C and Tim23 (**Figure 7C**). Similarly, *Bnip3*-KO is better than *Nix*-KO to rescue mitochondrial protein levels in *Fbxl4^−/−^* heart (**Figure S5A**). But interestingly, both *Bnip3*-KO and *Nix*-KO rescued mitochondrial proteins to similar levels in *Fbxl4^−/−^* kidney (**Figure S5B**). For similar reason as **Figure 6C**, we did not show the high molecular weight region of BNIP3 and NIX blots in **Figures 7C**, **S5A** and **S5B**. Collectively, these results suggest that *Bnip3* and *Nix* have variable contributions to mitophagy in different tissues/organs of the *Fbxl4^−/−^* mice.

Finally, we analyzed P0 liver metabolites to understand the effect of reduced mitochondrial function on cellular metabolism. Cellular ATP level was not altered in *Fbxl4^−/−^* liver, indicating bioenergetics is largely normal. We observed that pyruvate, lactate and several amino acids alanine, proline, tyrosine, isoleucine and aspartate greatly accumulated in *Fbxl4^−/−^* liver (**Figure 7D**), likely due to reduced oxidation of these metabolites in mitochondria. *Bnip3*-KO and *Nix*-KO, particularly the former, rescued these metabolic derangements in *Fbxl4^−/−^* liver (**Figure 7D**).

## DISCUSSION

The regulatory mechanisms of mitophagy remain poorly characterized. Here we reveal that FBXL4 forms a mitochondrial SCF-FBXL4 ubiquitin E3 ligase complex that ubiquitinates and degrades mitophagy receptors BNIP3 and NIX to control mitophagy at basal conditions. It is noteworthy that BNIP3 and NIX are responsive to diverse signals to regulate mitophagy. For example, BNIP3 and NIX are transcriptionally induced by HIF1α under hypoxia and in tumors (Bellot, Garcia-Medina et al., 2009, Vara-Pérez, Rossi et al., 2021, Zhang et al., 2008), NIX is induced during erythrocyte maturation (Sandoval et al., 2008, Schweers et al., 2007), and BNIP3 is induce by FoxO3 during muscle atrophy (Mammucari, Milan et al., 2007). Therefore, FBXL4 may regulate mitophagy under diverse developmental and stress/pathological conditions.

The detailed molecular mechanisms of SCF-FBXL4 complex assembly and substrate recognition remain unclear. It is reasonable to predict that some FBXL4 pathogenic mutations may disrupt complex assembly, and others may interfere with substrate binding. We were surprised to find that all the tested FBXL4 mutants (V140A, L481P, R482W, I551N, D565G, I566M and Q597P) impair complex assembly (**Figure 5G**). Testing more FBXL4 pathogenic mutations in future studies may reveal other types of pathogenic mechanisms. Another important question is that whether BNIP3 and NIX degradation by SCF-FBXL4 is a constitutive or regulated process. BNIP3 and NIX are subjected to post-translational modifications, such as phosphorylation (Onishi et al., 2021). Under hypoxia, phosphorylation of BNIP3 at Ser 60/Thr 66 by JNK1/2 was suggested to inhibit proteasomal degradation of BNIP3 and promote mitophagy (He, Gong et al., 2020). It is interesting to examine whether phosphorylation of BNIP3 and NIX regulates their degradation by SCF-FBXL4.

FBXL4 mutations cause severe multisystem degeneration in human patients (Bonnen et al., 2013, Gai et al., 2013) and in knockout mice (Alsina et al., 2020). The underlying cause has been unclear. Here we provide clear *in vivo* evidence that hyperactive mitophagy mediated by BNIP3 and NIX causes lethality of the *Fbxl4^−/−^* mice (**Figures 6** and **7**). Because FBXL4 pathogenic mutants are defective in degrading BNIP3 and NIX (**Figure 5**), it is highly likely that hyperactive mitophagy causes pathogenesis and lethality in human patients. Based on our results, rational design of therapeutic strategies can be conceived. For example, patients may benefit from chemical degraders of BNIP3 and NIX and from genetic approaches silencing BNIP3 and NIX, such as gene editing, RNAi, and anti-sense oligos etc. On the other hand, pharmacological approaches to promote mitochondrial biogenesis via targeting the PGC1 coactivators (Finck & Kelly, 2006, Spiegelman, 2007, Zhang, Zhou et al., 2013) may also be helpful. Correcting the metabolic defects of FBXL4-related patients is another angle to tackle the disease. Pyruvate is among the top metabolites that accumulate in the *Fbxl4^−/−^* mice (**Figure 7D**), indicating insufficient pyruvate catabolism may contribute to disease pathogenesis. Consistent with this idea, chemical activation of pyruvate dehydrogenase by dichloroacetate (DCA) improves symptoms in FBXL4-mutant worm and zebrafish (Lavorato, Nakamaru-Ogiso et al., 2022).

Mitophagy is an essential mitochondrial quality control mechanism. Impairment of mitophagy compromises stress resistance and diminishes lifespan extension by low insulin/IGF-1 signaling (Palikaras, Lionaki et al., 2015). Mitophagy impairment is also implicated in Parkinson’s disease (Pickrell & Youle, 2015). From this perspective, mitophagy activators have been actively sought and tested in animal models and human subjects for beneficiary effects (Andreux, Blanco-Bose et al., 2019, Hertz, Berthet et al., 2013, Ryu, Mouchiroud et al., 2016). Our results highlight that mitophagy activation has an upper limit and surpassing this limit may cause tissue/organ degeneration. Indeed, under stress conditions, cells/animals transcriptionally activate mitophagy and mitochondrial biogenesis together to accelerate mitochondrial removal while maintaining mitochondrial mass (Liu, Li et al., 2021a, Palikaras et al., 2015). Hence, coordinated activation of mitophagy and mitochondrial biogenesis may be a better strategy to improve mitochondrial quality control and maintain mitochondrial function.

## Materials and Methods

### Antibodies

BNIP3 (Abcam, ab109362; Abcam, ab10433; Cell signaling, 3769S), NIX (Cell signaling, 12396S), FUNDC1 (From Quan Chen lab), FLAG (Sigma-Aldrich, F1804), HA (Sigma-Aldrich, H6533), ACTIN (Sigma-Aldrich, A2066), TOM70 (Proteintech, 14528-1-AP), TOM20 (ABclonal, A19403), SMAC (Cell signaling, 2954S), MITOFILIN (Proteintech, 10179-1-AP), HSP60 (Cell signaling, 4870S), TIMM23(Proteintech, 11123-1-AP), SKP1 (Cell signaling, 2156S), CUL1 (Proteintech, 12895-1-AP), Total OXPHOS Cocktail (Abcam, ab110411), Cytochrome C (BD Biosciences, 556433), mCherry (Easybio, BE2026), MTCH2 (Proteintech, 16888-1-AP), BECLIN1 (Cell signaling, 3495T), FIP200 (ABclonal, A14685), TFAM (Abcam, ab131607), LONP1 (Proteintech, 15440-1-AP), VDAC (Cell signaling, 4866S), Donkey polyclonal anti-Rabbit IgG (H+L),HRP-conjugated (Jackson ImmunoResearch, 711-035-152), Donkey polyclonal anti-Mouse IgG (H+L), HRP-conjugated (Jackson ImmunoResearch, 711-035-151), Goat polyclonal anti-Mouse IgG, Fcγ fragment specific, HRP-conjugated (Jackson ImmunoResearch, 115-035-008), Goat polyclonal anti-Mouse IgG, light chain specific, HRP-conjugated (Jackson ImmunoResearch, 115-035-174).

### Chemicals and Reagents

Sodium pyruvate (Sigma-Aldrich, P5280), Uridine (Sigma-Aldrich, U3003), Doxycycline hyclate (Dox, Sigma-Aldrich, D9891), Polybrene (Sigma-Aldrich, 107689), Polyethylenimine (Polysciences, 23966-2), Opti-MEM™ (GIBCO, 31985070), Dulbecco’s modified Eagle’s medium (DMEM, GIBCO, C11965500BT), Fetal bovine serum (FBS, GIBCO, 10091148; Gemini, 900-108), Penicillin and streptomycin (Pen Strep, GIBCO, 15140122), Puromycin (InvivoGen, ant-pr-1), Blasticidin (InvivoGen, ant-bl-1), Geneticin (G-418, Amresco, e859-5), Hygromycin B (Sigma-Aldrich, 31282-0409), PhosSTOP (phosphatase inhibitor, Roche, 4906837001), cOmplete™, EDTA-free Protease Inhibitor Cocktail (Roche, 4693132001), Phenylmethanesulfonyl fluoride (PMSF, Sigma-Aldrich, P7626), Dithiobis (succinimidyl propionate) (DSP, Thermo Scientific, 22585).

### Recombinant DNA

Recombinant DNA used and generated in this paper were listed in **Table S4**.

### Cell lines and cell culture

HeLa and HEK293T were cultured in DMEM supplemented with 10% Fetal Bovine Serum and 1% penicillin-streptomycin. Both cell lines were incubated in tissue culture incubators at 37°C with 5% CO_2_.

### Mice

Rosa26-Cas9 knockin mice were purchased from The Jackson Laboratory. *Fbxl4^−/−^* mice and *Bnip3^−/−^* mice were purchased from GemPharmatech (Nanjing, Jiangsu, China). *Nix^−/−^* mice were generated by CRISPR-Cas9 genome editing technology and embryonic microinjection (sgRNA sequences in **Table S5**). *Fbxl4^+/-^* mice were crossed with *Bnip3^−/−^* mice and *Nix^−/−^* mice respectively to generate the *Fbxl4^−/−^Bnip3^−/−^* mice and *Fbxl4^−/−^Nix^−/−^* mice. Animals were maintained under a 12-hours light/dark cycle and on a standard chow diet at the specific pathogen-free (SPF) facility at the National Institute of Biological Sciences, Beijing. Mice were weighed at 30 days of age. All mouse experiments were carried out following the national guidelines for housing and care of laboratory animals (Ministry of Health, China) and performed in accordance with institutional regulations after review and approval by the Institutional Animal Care and Use Committee at National Institute of Biological Sciences, Beijing.

### Lentivirus production and generation of stable cell lines

Lentivirus was packaged by transfecting plasmids and lentiviral packaging vectors into HEK293T cells with 70% confluency using polyethylenimine (PEI) at a ratio of 1 μg plasmid: 5 μl PEI (1mg/ml). For each well of a six-well plate, a total of 2 µg plasmids (plasmids: psPAX2: pMD2.G = 5:3:2) was used. 48 hours later, the supernatants were collected and filtered through a 0.45 μm filter. Targeted cells were infected with lentivirus in the presence of 8 μg/ml polybrene for 48 h. Infected cells were selected by 2 μg/ml puromycin (for FUIPW vector), 20 μg/ml blasticidin (for plenti-Tet-On vector), 2.5 mg/ml G418 (for pLVX vector) or 200 μg/ml hygromycin (for plenti-Hygro vector) supplemented medium for 4 days. Gene expression was validated by immunoblotting.

### Generation of kncokout and knockin cell lines

The sequences of gRNAs are listed in **Table S5**. For single knockout clones, annealed gRNAs of target genes were ligated into the pX458 vector. The plasmids were transiently transfected into HeLa cells using PEI as the transfection reagent. 2 days later, GFP-positive cells were sorted into 96-well plate (one cell per well) by flow cytometry with the BD FACSAria Fusion. Single clones were grown for 2-3 weeks, and the resultant colonies were expanded and examined by immunoblotting and sequencing. For pool knockout, annealed gRNAs were ligated into the lentiCRISPR v2 vector. The plasmids were used to package lentivirus to infect the targeted cells as described above. Infected cells were selected by 2 μg/ml puromycin for 48 hours and the knockout efficiency was determined by immunoblotting. For FLAG knockin cells, annealed gRNAs of target genes were also cloned into the pX458 vector. The HMEJ donor containing the FLAG sequence flanked by 800-bp homology arms complementary to the knockin site with PAM mutation was inserted into pBM16A T-vector (Yao, Wang et al., 2017). For each well of a six-well plate, a total of 2 µg plasmids (pX458: donor = 1:1) was used. After 48 hours, GFP-positive HeLa cells were sorted into 96-well plates (one cell per well) by flow cytometry with the BD FACSAria Fusion. FLAG knockin clones were examined by immunoblotting and confirmed by DNA sequencing.

### CRISPR knockout screen

Lentiviral human mitoCarta2.0 CRISPR knockout library was generated as described (**He et al., Cell Reports, accepted**). A total of 5×10 ^6^ HEK293T cells were plated in 10-cm dishes. The following day, 10 µg of plasmids containing the sgRNA library, psPAX2, and pMD2.G at a mass ratio of 5:3:2 were transfected into HEK293T cells using the PEI transfection reagent. After 48 hours, lentivirus was harvested, aliquoted, and frozen at −80 °C. HeLa cells stably expressing cas9 and doxycycline-inducible mitoQC or mtkeima reporter was generated by lentiviral infection. The cas9-expressing reporter cells were infected with packaged sgRNA library at a multiplicity of infection (MOI) of approximately 0.3, keeping coverage of 1000 cells per sgRNA on average. After 24 hours of virus transduction, cells were selected for 7 days in the medium supplemented with 1 μg/ml puromycin, 50 μg/ml uridine and 3 mM pyruvate and maintained at 1000× coverage during each passage. Then the positively-transduced cells were treated with Dox for 48 hours followed by culturing in Dox-free medium for 24 hours. Cells were then collected for flow cytometry. Top 25% of cells with the strongest mitophagy and bottom 25% of cells with the lowest mitophagy were sorted (2.3 × 10^6^ cells for each sample).

Genomic DNA was extracted by Phenol/chloroform extraction. For the first step of sgRNA amplification, six PCRs (each containing 2.5 µg genomic DNA) with primer-OCY581-NGS-Lib-KO-Fwd-first-step and OCY582-NGS-Lib-KO-Rev-first-step was performed using NEBNext® Ultra™ II Q5® Master Mix (New England Biolabs, M0544L). The first-step PCR products were recovered with QIAquick Gel Extraction Kit (Qiagen, 28706) and used as DNA templates for six PCRs (each containing 2 ng purified first-step PCR products) with primer-OCY583-NGS-Lib-KO-Fwd-second-step and OCY (584-589)-NGS-Lib-KO-Rev-barcode. The second-step PCR products were separated by a 2% agarose gel, purified and sequenced by Nova-seq 150-bp paired-end sequencing (Anoroad, Beijing, China).

### Analysis of screen results

Raw reads were trimmed using Trim Galore v0.6.6 (https://www.bioinformatics.babraham.ac.uk/projects/trim_galore/) with default settings. The MAGeCK (0.5.9.3) was used to analyze the screening data. We used the MAGeCK “count” command to generate read counts and normalized using median normalization of all samples. The raw read counts of all sgRNAs for all samples were merged into a count matrix. We next used MAGeCK “test” command to identify the top negatively and positively selected sgRNAs or genes with default settings with software available from https://sourceforge.net/projects/mageck/.

### Mitophagy assay

HeLa cells stably expressing a doxycycline (Dox)-inducible mitoQC or mtKeima reporter were treated with 2 μg/ml Dox for 48 hours and subsequently without Dox for 24 hours unless otherwise indicated. After treatment, cells were trypsinized and filtered by a 40 μm filter for flow cytometry. Flow Cytometry was performed using the BD FACSAria Fusion with a BV605 detector for neutral pH and a PE-Texas Red detector for acidic pH. Data analysis was performed using FlowJo V10. 10,000 cells were analyzed per condition and all statistical analyses were performed using data from at least three biological replicates.

For live cell imaging of mtKeima, cells were cultured in 35 mm glass-bottom culture dish (NEST, 801002) and treated with Dox for 48 hours and subsequently without Dox for 24 hours. Samples were imaged without fixation by a NIKON A1 + SIM confocal microscope with a 60 × oil objective (CFI Plan Apochromat Lambda; NA1.40; Nikon). For mCherry cleavage assay, cells stably expressing a doxycycline-inducible mCherry-FIS1^101-152^ reporter were cultured in a 6-well plated and lysed for immunoblot after doxycycline treatment. The statistical analyses were performed using data from three biological replicates

### Immunofluorescence and confocal microscopy

HeLa cells were grown on glass coverslips and treated as described. After washing with PBS for 3 × 10 min, cells were fixed with 4% paraformaldehyde in PBS for 30 min at room temperature, permeabilized with 0.1% Triton X-100 in PBS for 30 min, blocked with 10% BSA in PBS for 30 min, and then stained with primary antibodies diluted in 10% BSA overnight at 4 °C. After washing with PBS for 3 × 10 min, cells were stained with Alexa-Fluor–conjugated secondary antibodies (1:500 dilution in PBS with 10% BSA) for 60 min at room temperature. Coverslips were then washed with PBS for 3 × 10 min and mounted onto slides with Fluoromount-G (SouthernBiotech, 0100-01). Fluorescent samples were examined with a NIKON A1 + SIM confocal microscope with a 60 × oil objective (CFI Plan Apochromat Lambda; NA1.40; Nikon).

### Immunoblotting

Cultured HeLa cells were washed once with PBS and scraped off to collect cell pellets. For mouse samples, neonatal mouse tissues were dissected out and snap-frozen in liquid nitrogen. Tissues were weighted and homogenized with the tissue homogenizer (DHS life science, Q24RC) at 6.5 m/s for 1 min at 4 °C. Homogenization was performed for two times with 1 min interval on ice. Both types of samples were lysed for 30 min on ice with FLAG Lysis Buffer (50 mM Tris–HCl pH 8.0, 1 mM EDTA, 1% Triton X-100, 150 mM NaCl, 10% Glycerol, 10 mM NaF) supplemented with 1 × cOmplete, EDTA-free protease inhibitor cocktail and 1 × PhosSTOP phosphatase inhibitor cocktail. The lysates were then centrifuged at 15,000 rpm for 10 min at 4 °C to collect the supernatant. Protein concentrations were determined by Bradford assay kit (Sigma-Aldrich, B6916). Lysates were mixed with 5 × sample buffer (250 mM Tris-HCl, pH 6.8, 50% glycerol, 5% 2-mercaptoethanol, 10% SDS, 0.08% Bromophenol Blue) followed by boiling at 98 °C for 15 min. Protein samples (5 ~30 µg in amount) were separated on SDS–PAGE gels and transferred to PVDF membranes. Membranes were incubated with primary antibodies in 5% nonfat milk in PBST overnight at 4 °C. Membranes were washed three times in PBST for 5 min once after blotting with primary antibodies. Then, the membranes were incubated with corresponding secondary HRP-conjugated antibodies for 1 hour at room temperature. Membranes were washed three times in PBST before visualization with ECL reagents.

### qPCR

HeLa cells were cultured in 6-cm dishes and treated as described. Cells were then lysed in 1 ml TRIzol reagent (Invitrogen, 15596026) and incubated at room temperature for 5 minutes. 0.2 ml chloroform was added and fully mixed. After incubating for 2 min, samples were centrifuged at 12, 000 g for 15 minutes at 4 °C. Then, 500 μl RNA-containing aqueous phase was transferred to a new tube and another 500 μl isopropanol was added to it. The mixture was incubated at −20 °C for 6 min and at room temperature for 10 minutes. RNA pellet was collected by centrifuging at 12,000 g for 10 minutes at 4 °C and washed with 75% ethanol. The RNA pellet w as air-dried for 10 min and resuspended in nuclease-free H_2_O. RNA was converted into cDNA using 5x ALL-In-One RT Master Mix (Applied Biological Materials Inc, G490) following the manufacturers’ instructions and diluted 20 times with nuclease-free H_2_O. Quantitative PCR was performed using Taq Pro Universal SYBR qPCR Master Mix (Vazyme, Q712-03) in a CFX96 Touch Real-Time PCR Detection System (Bio-Rad) with 3 biological replicates. The relative gene expression levels were normalized to ACTB. Primers used for qPCR were listed in **Table S5**.

### Submitochondrial localization analysis

HeLa cells cultured in six 15-cm dishes were scraped to collect cell pellets. Cell pellets were then resuspended with 7 ml homogenization buffer (20 mM HEPES/KOH pH=7.4, 220 mM mannitol, 70 mM sucrose, 1 mM EDTA, 0.5% (W/V) BSA) supplemented with 1 mM PMSF and protease inhibitor cocktail and incubated for 20 min on ice. Cells were homogenized by a Teflon potter (Glas-Col) rotating at 600 rpm by moving the pestle up and down for 5 repeats. Cell homogenates were centrifuged at 1,000 g for 5 min at 4 °C. This step was repeated at least three times to completely discard cell debris and the nuclear fraction. Supernatants were divided into seven aliquots and subsequently transferred to 1.5 ml Eppendorf tube, followed by centrifugation at 18,000 g for 10 min at 4 °C to pellet mitochondria. The mitochondrial pellets were washed by resuspending the pellet with 1 ml homogenization buffer and centrifuged again to wash out the protease inhibitors.

For six mitochondrial aliquots, two were resuspended in 300 μl homogenization buffer, two resuspended treated with 300 μl hypotonic swelling buffer (10 mM HEPES/KOH pH 7.4, 1 mM EDTA), and two resuspended with 300 μl homogenization buffer supplemented with 0.5% (V/V) Triton X-100. One aliquot of each treatment was treated with proteinase K (66.7 μg/ml) for 20 min on ice while the other aliquot was left untreated as control. The seventh aliquot was resuspended with homogenization buffer as above, followed by treatment with trypsin (25 ng/ml) for 30 min at room temperature. The mitochondrial proteins were then precipitated by 300 μl 30% TCA (W/V) and incubated on ice for 10min. The mitochondrial proteins were collected by centrifuging at 18,000 g for 10 min at 4 °C. The protein pellets were washed with 1 ml 100% ethanol and centrifuged again. The pellets were dissolved in 100 μl sample buffer (60 mM Tris–HCl, pH 6.8, 7% glycerol, 2% 2-mercaptoethanol, 2% SDS, 0.02% Bromophenol Blue) and boiled at 98 °C for 10 min. 10 μl sample was used for immunoblotting.

### Analysis of the membrane association of proteins

Mitochondria were isolated as described above and divided into five aliquots. Mitochondrial pellets were resuspended in 800 μl freshly prepared Na_2_CO3 (100 mM) with pH values of 10, 11, 11.5, 12, and 12.5 respectively, and incubated for 30min on ice. The soluble and insoluble fractions were separated by centrifugation at 100,000 g at 4 °C for 1 hour. The supernatant soluble fractions were precipitated by 800 μl TCA and dissolved in 70 μl sample buffer as described above. The insoluble pellets were dissolved in 100 μl sample buffer and boiled at 98 °C for 10 min. 10 μl sample was used for immunoblotting.

### Immunoprecipitation

HeLa cells expressing FLAG-tagged baits were cultured in a 10-cm dish, washed once with PBS, and scraped to collect cell pellets. Cell pellets were lysed by 1 ml FLAG Lysis Buffer supplemented with 1 mM PMSF, 1 × protease inhibitor cocktail and 1 × phosphatase inhibitor for 30 min on ice. Cell extracts were centrifuged at 15,000 g for 10 min at 4 °C to remove cell debris. Supernatants were incubated with 8 µL anti-FLAG agarose beads (Sigma-Aldrich, A2220) for 8 hours at 4 °C. The beads were washed five times with the FLAG lysis buffer and eluted with 60 µL FLAG lysis buffer supplemented with 2 mg/ml FLAG peptide (ChinaPeptides), 1 × protease inhibitor cocktail and 1 × phosphatase inhibitor overnight at 4 °C. The eluted products were boiled at 98 °C for 10 min and analyzed by immunoblotting.

For analyzing the interaction between FBXL4-3FLAG and UBXD8-VCP, buffer A (50 mM HEPES-KOH (pH 7.5), 50 mM Mg (OAc)2, 70 mM KOAc, 0.2% Triton X-100, 0.2 mM EDTA, 10% glycerol) was used instead of the FLAG lysis buffer.

### DSP crosslinking and LC-MS analysis of proteins

FBXL4-KO HeLa cells stably expressing WT, V140A or I551N FBXL4-FLAG were cultured in twelve 15-cm dishes respectively. Collected cell pellets were resuspended with 6 ml DSP Lysis Buffer (40 mM HEPES, pH 7.5, 120 mM NaCl, 1 mM EDTA, 1% Triton X-100) supplemented with 2 mM DSP (Thermo Scientific, 22585), 2 mM ATP, 1 mM PMSF, 1 × protease inhibitor cocktail and 1 × phosphatase inhibitor for 30 min on ice. The lysates were diluted with 18 ml DSP Lysis Buffer supplemented with 1 × protease inhibitor cocktail, 1 × phosphatase inhibitor and 2mM ATP. Crosslinking reactions were quenched by adding 6 ml 1 M Tris-HCl, pH 7.4 followed by additional 30 min incubation on ice. Cell extracts were centrifuged at 15,000 g for 10 min at 4°C to remove cell debris. Supernatants were incubated with 100 µL anti-FLAG agarose beads for 8 hours at 4°C. The beads were washed by rotating at 4 °C for 5 minutes with DSP Lysis Buffer for the first time, DSP Lysis Buffer supplemented with 0.5M NaCl for two times, and DSP Lysis Buffer for two additional times. The beads were eluted with 80 µL DSP Lysis Buffer supplemented with 2 mg/ml FLAG peptide, 1 × protease inhibitor cocktail and 1 × phosphatase inhibitor overnight at 4 °C. The eluted products were boiled at 98 °C for 10 min.

Protein samples were separated by SDS-PAGE (migrating into the separation gel for 2 cm). The SDS-PAGE gels were stained by Coomassie Blue G250 overnight at room temperature and de-stained with H_2_O on the second day. The 2-cm gel containing all the proteins were cut and in-gel digested with sequencing grade trypsin (10 ng/μl trypsin, 50 mM ammonium bicarbonate, pH 8.0) overnight at 37 °C. Peptides were extracted with 5% formic acid/50% acetonitrile and 0.1% formic acid/75% acetonitrile sequentially and then concentrated to ~ 20 μl. The extracted peptides were separated by an pre-column (100 μm × 2 cm) packed with 3 μm spherical C18 reversed phase material (Dr.maisch, GmbH, Germany) and analytical capillary column (100 μm × 15 cm) packed with 1.9 μm spherical C18 reversed phase material (Dr.maisch, GmbH, Germany). A Waters nanoAcquity UPLC system (Waters, Milford, USA) was used to generate the following HPLC gradient: 0-8% B in 10 min, 8-30% B in 30 min, 30-80% B in 15 min, 80% B in 5 min (A = 0.1% formic acid in water, B = 0.1% formic acid in acetonitrile). The eluted peptides were sprayed into a LTQ Orbitrap Velos mass spectrometer (ThermoFisher Scientific, San Jose, CA, USA) equipped with a nano-ESI ion source. The mass spectrometer was operated in data-dependent mode with one MS scan followed by ten HCD (High-energy Collisional Dissociation) MS/MS scans for each cycle. Database searches were performed on an in-house Mascot server (Matrix Science Ltd., London, UK) for proteins mass spectrometric analysis. The search parameters are: 10 ppm mass tolerance for precursor ions; 0.1 Da mass tolerance for product ions; two missed cleavage sites were allowed for trypsin digestion and the following variable modifications were included: oxidation on methionine, Acetyl (Protein N-term).

### Adeno-associated virus (AAV) packaging and hepatic mitophagy measurement

The AAV packaging and injection method was described previously (Liu et al., 2021b) with slight modification. Non-targeted control (NTC) gRNA or a tandem cassette of 3 gRNAs targeting FBXL4 was cloned into AAV-U6-gRNA-CAG-mtKeima-WPRE-hGHpA (modified from Addgene, 60229) under the U6 promoter. The gRNA sequences were listed in **Table S4**. AAV-pro 293T cells were transfected with gRNA vectors along with AAV packaging vectors AAV2/9 and pAdDeltaF6 using the PEI MAX transfection reagent (Polysciences, 24765). 72 hours after transfection, cell pellets were harvested by cell lifter (Biologix, 70-2180) and suspended in 1× Gradient Buffer (10mM Tris-HCl pH=7.6, 150mM NaCl, 10mM MgCl2). Cells were lysed by five repeated cycles of liquid nitrogen freezing, 37°C water bath thawing and vortex. Then, benzonase nuclease (50 U/ml, Sigma-Aldrich, E1014) was added to cell lysates and incubated at 37 °C for 30 minutes to eliminate cellular DNA. Centrifuge the cell lysate at 21,130g for 30 min at 4°C and carefully transfer the supernatant to a pre-build iodixanol step gradients (15%, 25%, 40% and 58%, Sigma-Aldrich, D1556)) for ultracentrifugation purification at 41,000 rpm for 4 hours, 4°C. Accurately insert the needle ~1-2 mm below the interface between the 40% and 58% gradient and extract all the 40% virus containing layer. Purified AAV9 were concentrated using Amicon filters (EMD, UFC801096) and formulated in phosphate-buffered saline (PBS) supplemented with 0.01% Pluronic F68 (GIBCO, 24040032). Virus titers were determined by qPCR using a linearized AAV plasmid as a standard.

At P35, Rosa26-Cas9 knockin mice were randomly assigned to receive an intravenous injection of 200 µL of the NTC or FBXL4-targeting AAV9 virus (1.5×10^10^ genome copies/µL) through retro-orbital injection. 45 days after AAV injection, mice were sacrificed. Mouse livers were collected in multiple aliquotes. One aliquote was immediately washed with PBS, placed into 35 mm glass-bottom culture dish containing PBS and examined with a NIKON A1 + SIM confocal microscope with a 60 × oil objective. The other aliquotes were snap-frozen in liquid nitrogen and stored at −80 °C for immunoblotting.

### Extraction and LC-MS Analysis of metabolites from mouse livers

P0 mouse livers were dissected out and snap-frozen in liquid nitrogen. Livers were weighted and resuspended with ice-cold acetonitrile (Sigma-Aldrich, 34851)/methanol (Sigma-Aldrich, 34860)/water (4/4/2) (20 μL/mg). Tissues were homogenized at 6.5 m/s for 1 min at 4 °C in the tissue homogenizer (DHS life science, Q24RC) followed by ice incubation for 2 min. Repeat homogenization and cooling for 2 times. Equal volume of tissue homogenates was transferred to new Eppendorf tubes. Tissue homogenates were centrifuged at 20,000 g for 10 min at 4°C and the supernatants were transferred to new Eppendorf tubes. The supernatants were dried by a Speedvac vacuum concentrator at 4 °C. The dried samples were stored at −80 °C and analyzed within 24 h. The remaining pellets were dissolved in 0.1 M KOH by shaking at 4 °C overnight. Protein concentration was measured by Bradford. The metabolic data was normalized to protein concentration.

Authentic reference standard compounds were purchase from Sigma-Aldrich (St. Louis, MO, USA). Stock solutions of 1 mg/ml were prepared in water. Working solutions of 10, 20, 50, 100, 200, 500, 1000, 2000, 5000, 10000 ng/ml were obtained by serial dilution of stock solutions in methanol/water (1:1). Samples were resuspended in 100 μl 50% methanol and filtered through 0.45 μm filters. The standard solutions and biological extracts were transferred to 250 μl inserts in auto-sampler vials and 6 µl of each sample was injected to LC-MS.

The LC-MS analysis was performed using a Thermo Vanquish UHPLC coupled to a Thermo Q Exactive HF-X hybrid quadrupole-Orbitrap mass spectrometer. A Merck ZIC-cHILIC column (2.1×100 mm, 3 μm) was used for separation. The mobile phases consisted of 10 mM ammonium acetate in 5/95 ACN/water (A) and 10 mM ammonium acetate in 95/5 ACN/water (B). The following gradient was applied: 0-5 min, 99% B; 5-20 min, 99-20% B; 20-21 min, 20-99% B; 21-25 min, 99% B. The flow rate was 0.5 ml/min, and the column temperature was 40 °C. Full-scan mass spectra were acquired in the range of m/z 66.7 to 1000 with the following ESI source settings: spray voltage 3.5 kV, aux gas heater temperature 380 °C, capillary temperature 320 °C, sheath gas flow rate 35 units, aux gas flow gas 10 units in the positive mode, and spray voltage 2.5 kV, aux gas heater temperature 380 °C, capillary temperature 320 °C, sheath gas flow rate 30 units, aux gas flow gas 10 units in the negative mode. MS1 scan parameters included resolution 60000, AGC target 3e6, and maximum injection time 200 ms. The results were analyzed using Thermo Scientific Xcalibur software.

### Quatification

For quantifying mCherry cleavage percentage, the intensity of full-length and cleaved mCherry immunoblotting bands was measured with the ImageJ software. For measuring mitolysosome area/mitochondria area, the area of mitolysosome and mitochondria were measured with the ImageJ software. Quantification results are represented as the mean ± standard deviation (SD).

### Statistical analyses

All the statistical details of experiments can be found in the figure legends. GraphPad Prism 8 was used to analyze the data. Data were presented as the mean + standard deviation (SD) unless otherwise indicated. Sample size (n) indicates biological replicates from a single representative experiment. For mouse study, sample size (n) indicates data from a distinct mouse. For mtKeima-based analysis of the percentage of cells with mitophagy, sample size (n) indicates the number of cells analyzed. For mtKeima-based analysis of mitolysosome area versus mitochondria area, sample size (n) represents the number of imaging areas analyzed in cell culture experiments and the number of cells analyzed in AAV experiments. The results of all experiments were validated by independent repetitions. For comparing of two groups, statistical significance was determined using a two-tailed unpaired Student’s t-test; For comparing multiple groups, statistical significance was determined by one-way ANOVA using Tukey-Kramer test. *P* values are denoted in figures as: not significant [ns], \**P* < 0.05, ** *P* < 0.01, *** *P* < 0.001.

## ACKNOWLEDGMENTS

We thank Dr. Chen Quan (Nankai University) and Dr. Liu Lei (Institute of Zoology, CAS) for the FUNDC1 antibody. This research was supported by the Beijing Municipal Science and Technology Commission and Tsinghua University.

## CONFLICT OF INTERESTS

The authors declare no competing interests

---

**Figure S1.**
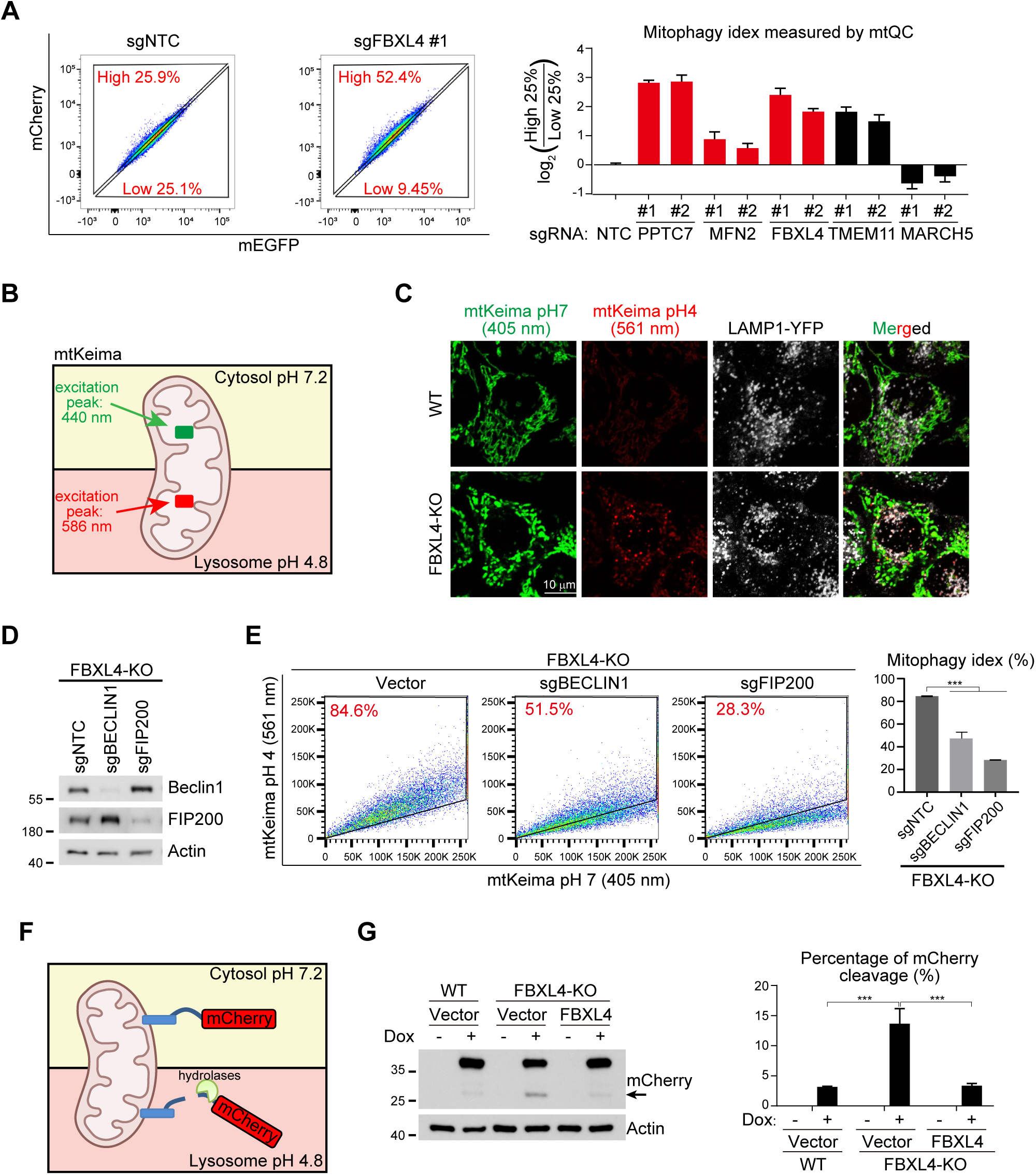
FBXL4 deficiency hyperactivates mitophagy. Supplemental to Figure 1. (A) Verification of screen hits by mitoQC-based FACS analysis of mitophagy. Left: representative FACS analysis; right: quantitative analysis of mitophagy levels in HeLa cells expressing the indicated sgRNAs. Two independent sgRNAs were used for each gene. NTC: non-targeting control. (B) Schematic of the pH-dependent dual excitation of mtKeima. (C) Live cell imaging analysis of mitolysosomes. WT and FBXL4-KO HeLa cells expressing mtKeima and LAMP1-YFP were subject to live cell confocal imaging. (D) Immunoblot analysis of the indicated HeLa cells. FBXL4-KO HeLa cells were infected with lentiviruses expressing sgNTC, sgBECLIN1 or sgFIP200. (E) FACS analysis of mitophagy levels in the indicated HeLa cells. The same cells in (D) were analyzed. Left: representative FACS analysis; right: quantitative analysis. (F) Schematic of a cleavage-based mitophagy reporter. The linker between mCherry and mitochondrial targeting sequence is sensitive to lysosomal hydrolases. (G) Immunoblot analysis of reporter cleavage in the indicated HeLa cells. Left: representative immunoblot. Arrow points to the cleaved mCherry. Right: quantitative analysis of mCherry cleavage. Data are mean + SD from three biological replicates (A, E, G). Statistics: two-tailed unpaired Student’s t-test (E, G); \*\**P* < 0.01; \*\*\**P* < 0.001.

**Figure S2.**
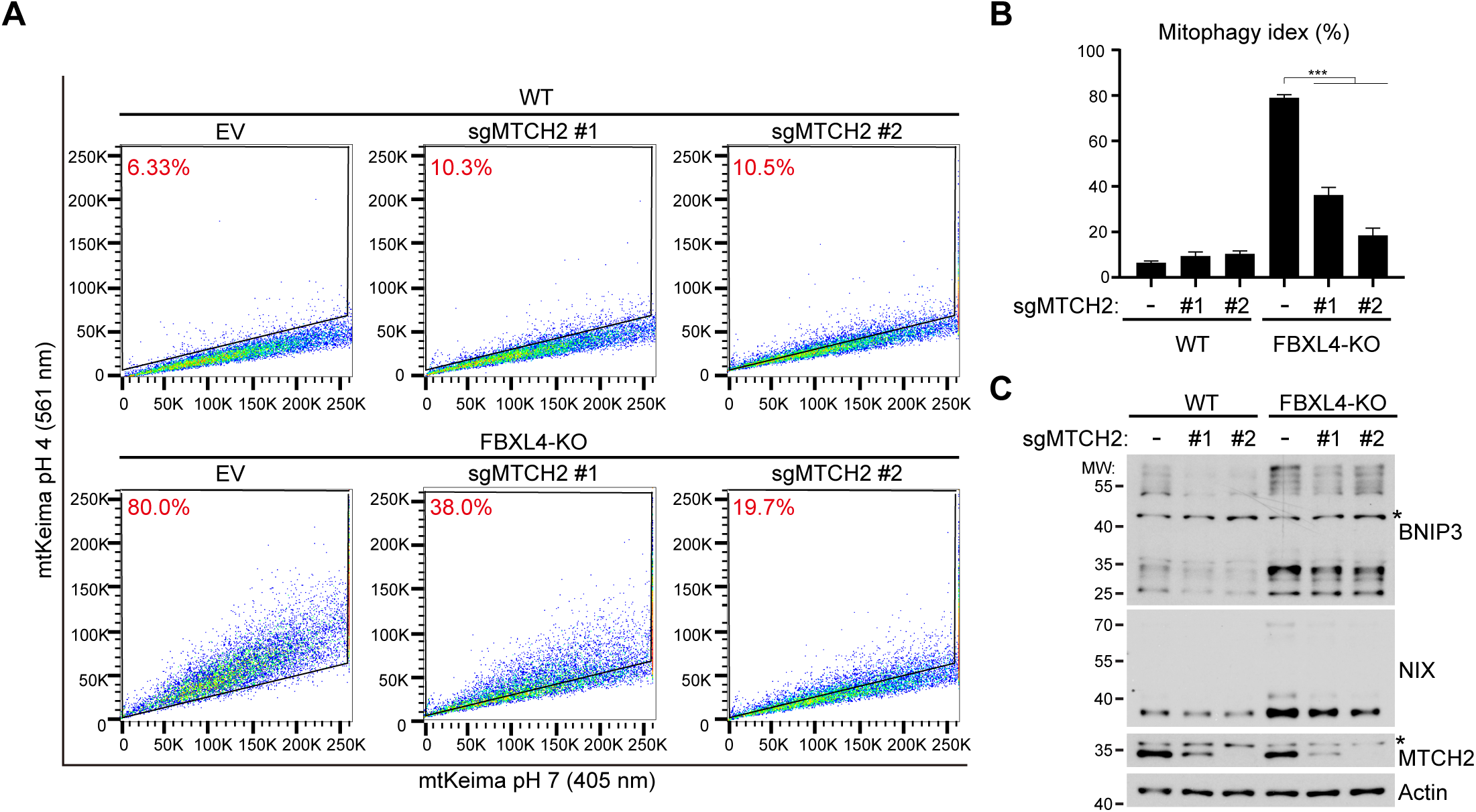
MTCH2 regulates the protein levels of BNIP3 and NIX and controls mitophagy in FBXL4-null cells. Supplemental to Figure 2. (A and B) Representative FACS analysis (A) and quantitative analysis (B) of mitophagy levels in the indicated HeLa cells. Two independent sgRNAs for MTCH2 were used. Data are mean + SD from three biological replicates. Statistics: two-tailed unpaired Student’s t-test; \*\*\**P* < 0.001. (C) Immunoblot analysis of the indicated HeLa cells. *: non-specific bands.

**Figure S3.**
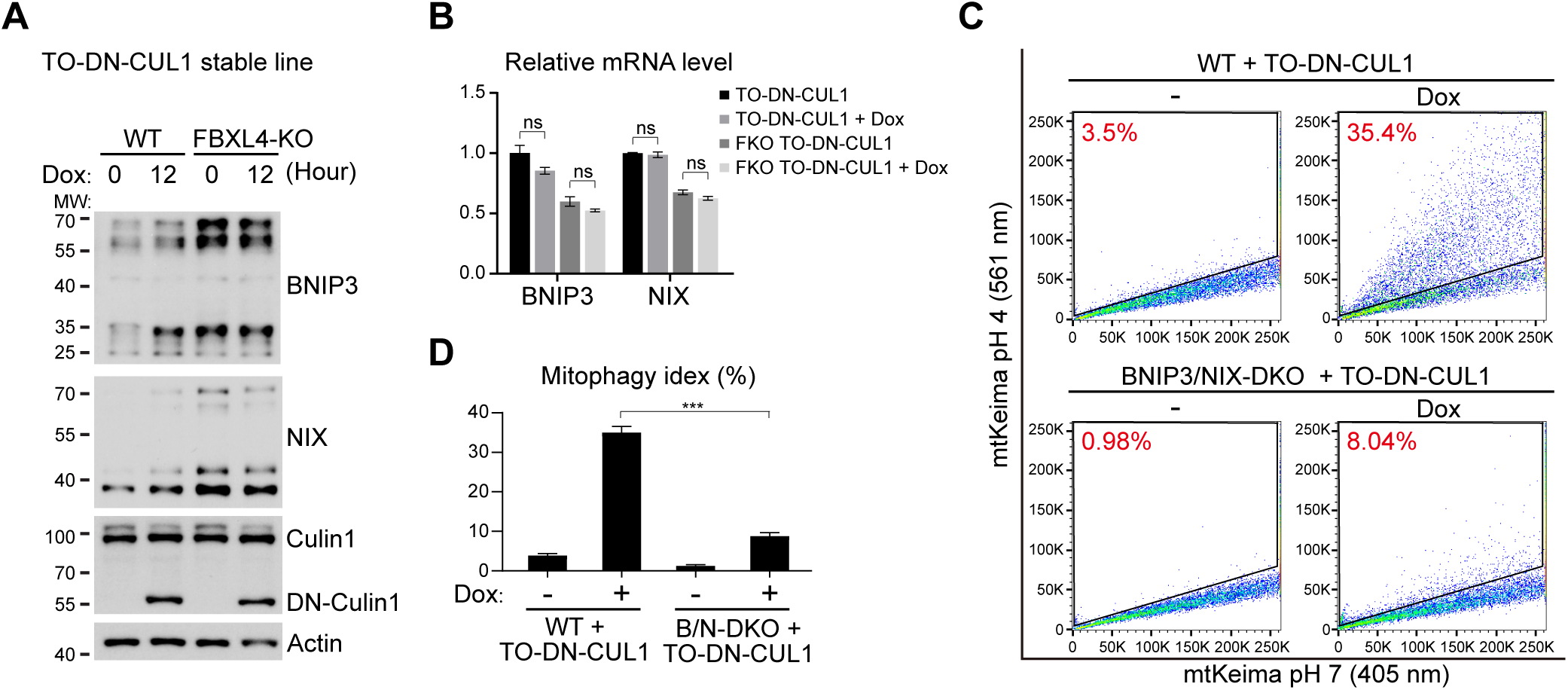
Overexpression of DN-Cullin1 causes the accumulation of BNIP3 and NIX to hyperactivate mitophagy. Supplemental to Figure 4. (A) Immunoblot analysis of the indicated HeLa cells. WT and FBXL4-KO HeLa cells were infected with lentivirus expressing TO-DN-Cullin1. Cells were then treated with doxycycline (Dox, 2 μg/ml) for the indicated time. (B) qPCR analysis of the indicated HeLa cells. The same cells in (A) were used. Dox (2 μg/ml) was treated for 12 hours. (C and D) Representative FACS analysis (C) and quantitative analysis (D) of mitophagy levels in the indicated HeLa cells. WT and FBXL4-KO HeLa cells were infected with lentivirus expressing TO-DN-Cullin1. Cells were then treated with Dox (2 μg/ml) for 48 hours. Data are mean + SD from three biological replicates (B, D). Statistics: two-tailed unpaired Student’s t-test (B, D); ns: not significant; \*\*\**P* < 0.001.

**Figure S4.**
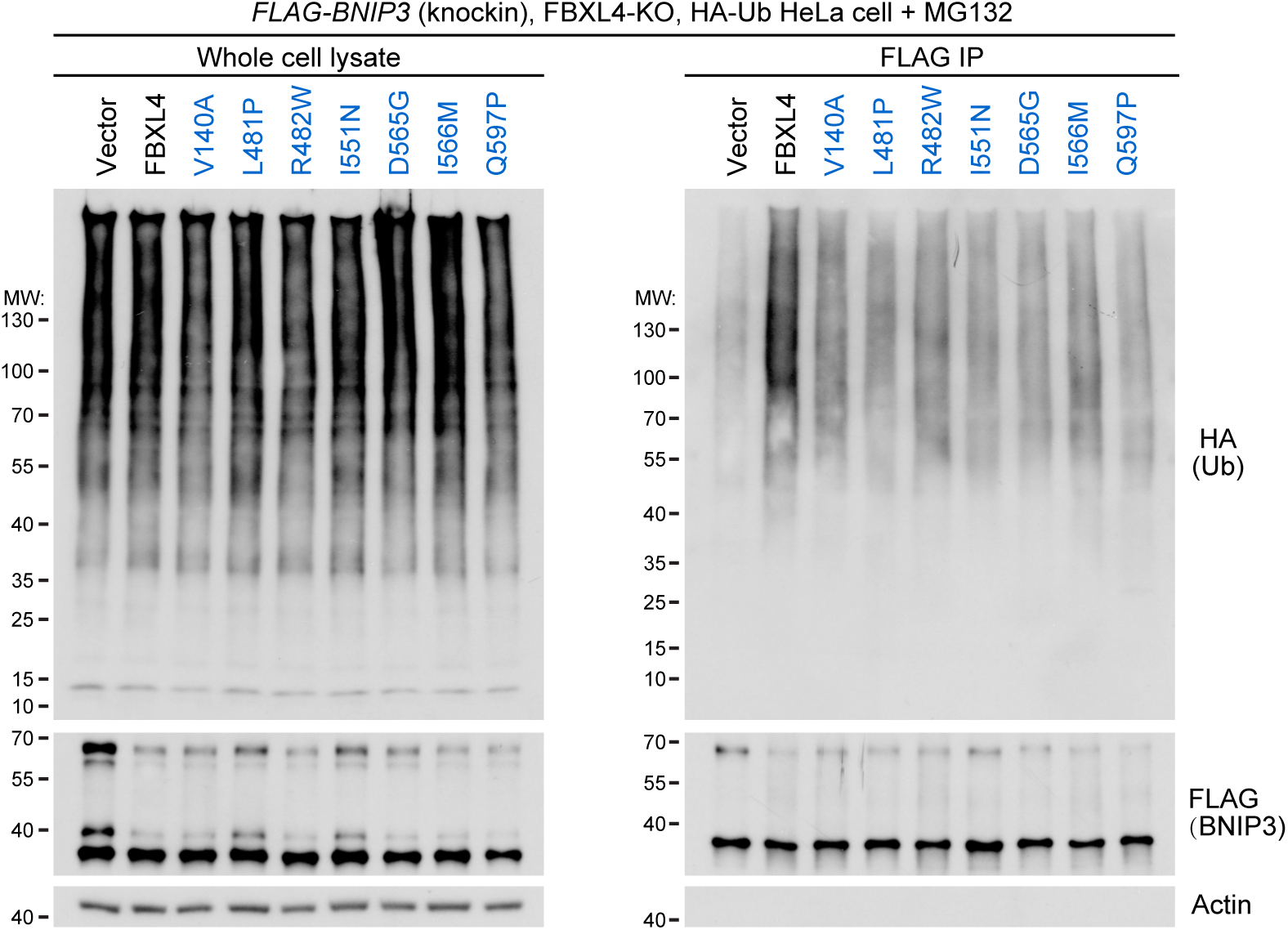
Pathogenic mutations of FBXL4 impair substrate ubiquitination. Supplemental to Figure 5. Immunoprecipitation analysis of the ubiquitination of BNIP3 by FBXL4 pathogenic mutants. FLAG-BNIP3 (knockin), FBXL4-KO, HA-Ub HeLa cells were rescued with WT or mutant FBXL4, and treated with DMSO or MG132 (20 μM) for 8 hours.

**Figure S5.**
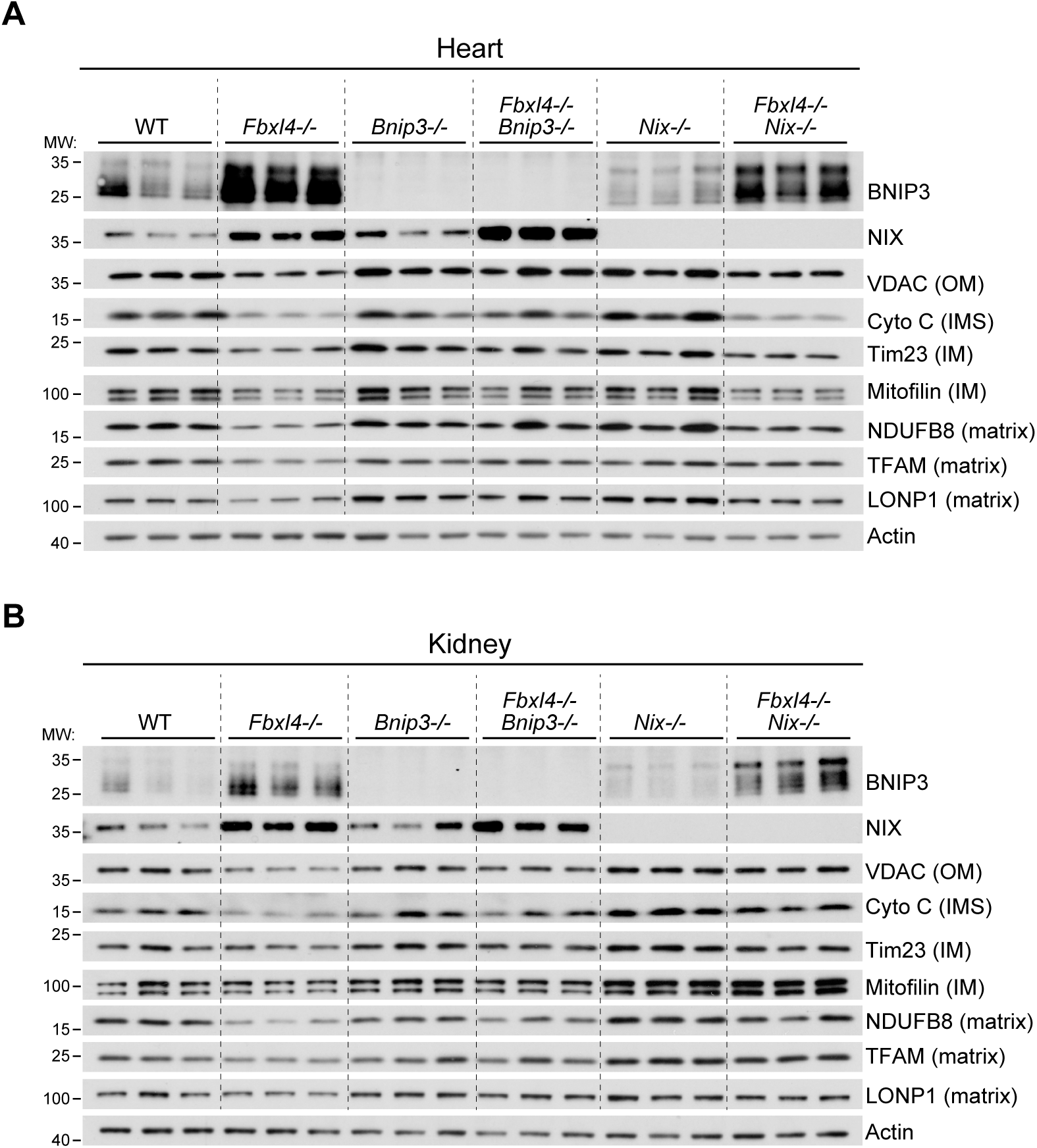
Tissue specific contribution of *Bnip3* and *Nix* to mitochondrial mass reduction in *Fbxl4^−/−^* mice. Supplemental to Figure 7. Immunoblot analysis of heart (A) and kidney (B) from the indicated mice at P0 of age. Three mice for each genotype were analyzed.

**Table S1. mitoQC-based mitophagy screen result of HeLa cells.**

**Table S2. mitoKeima-based mitophagy screen result of HeLa FBXL4-KO cells.**

**Table S3. Mass-spectrometry result of WT, V140A and I551N FBXL4-FLAG immunoprecipitation.**

**Table S4.** Recombinant DNA used in this study. Related to Methods.

| Recombinant DNA |  |  |
| --- | --- | --- |
| Plasmid: psPAX2 | Addgene | Cat# 12260 |
| Plasmid: pMD2.G | Addgene | Cat# 12259 |
| Plasmid: pSpCas9(BB)-2A-GFP (PX458) | Addgene | Cat# 48138 |
| Plasmid: lentiCas9-Blast | Addgene | Cat# 52962 |
| Plasmid: lentiGuide-Puro | Addgene | Cat# 52963 |
| Plasmid: lentiCRISPR v2 | Addgene | Cat# 52961 |
| Plasmid: AAV packaging vectors AAV9 | Addgene | Cat# 112865 |
| pAdDeltaF6 | Addgene | Cat# 112867 |
| Plasmid:<br>AAV-U6-nontargeted control-sgRNA-mtKeima | This paper | N/A |
| Plasmid: AAV-U6-sgFBXL4-mtKeima | This paper | N/A |
| Plasmid: plenti-Tet-On-mcherry-mEGFP-fis1 | This paper | N/A |
| Plasmid: plenti-Tet-On-mtKeima | This paper | N/A |
| Plasmid: plenti-Tet-On-mcherry-fis1-BSD | This paper | N/A |
| Plasmid: FUIPW | Feng Shao lab | N/A |
| Plasmid: FUIPW-FBXL4-3FLAG | This paper | N/A |
| Plasmid: pLVX-G418 | Ting Han lab | N/A |
| Plasmid: pLVX-FBXL4-3FLAG-G418 | This paper | N/A |
| Plasmid: plenti-EF1a-hgyro | Ting Han lab | N/A |
| Plasmid: plenti-EF1a-FBXL4-hgyro | This paper | N/A |
| Plasmid: plenti-Tet-On-CUL1-DN-3FLAG | This paper | N/A |
| Plasmid: FUIPW-LAMP1-YFP | This paper | N/A |
| Plasmid: FUIPW-3HA-Ub | This paper | N/A |

**Table S5.**
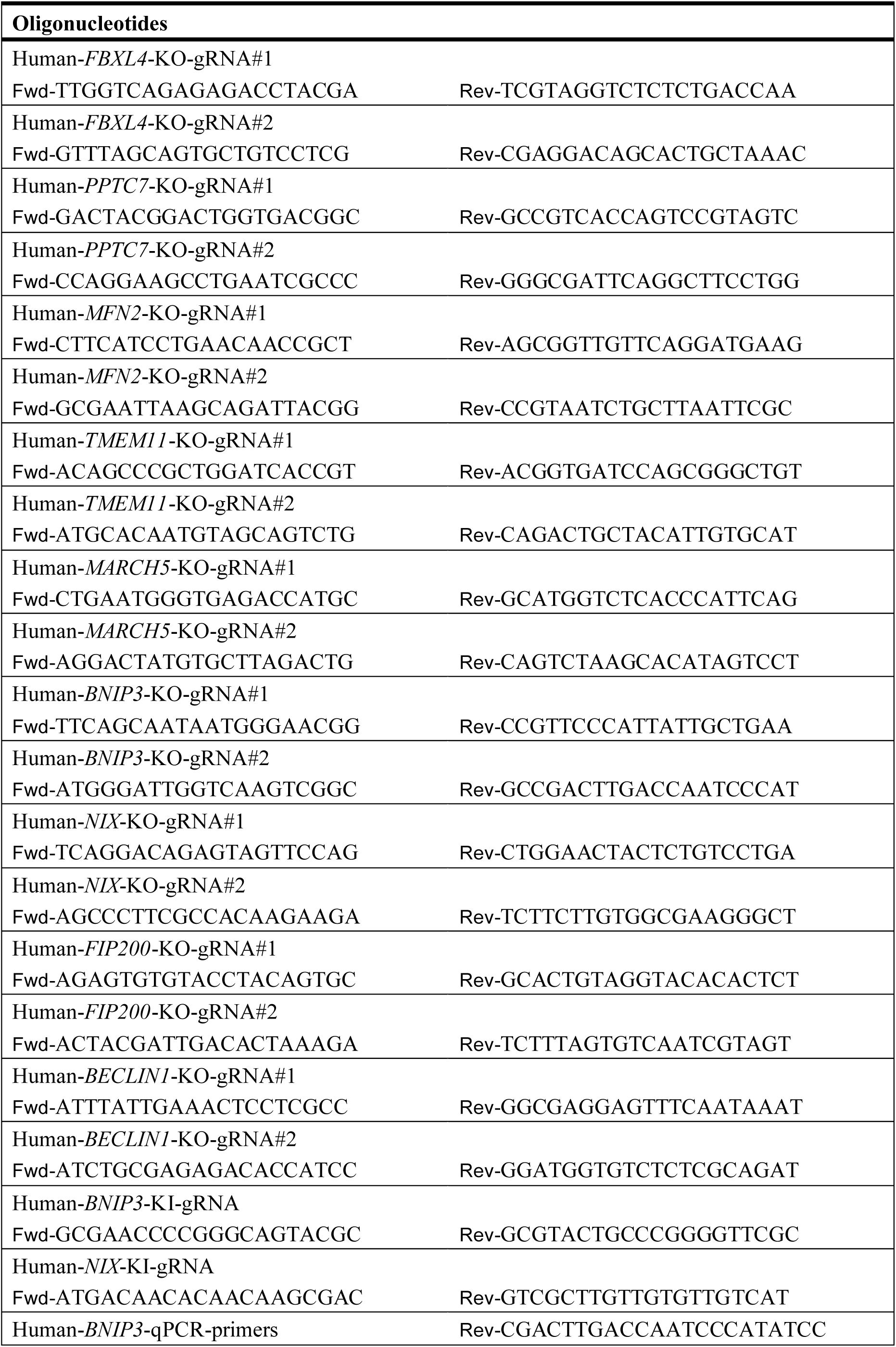
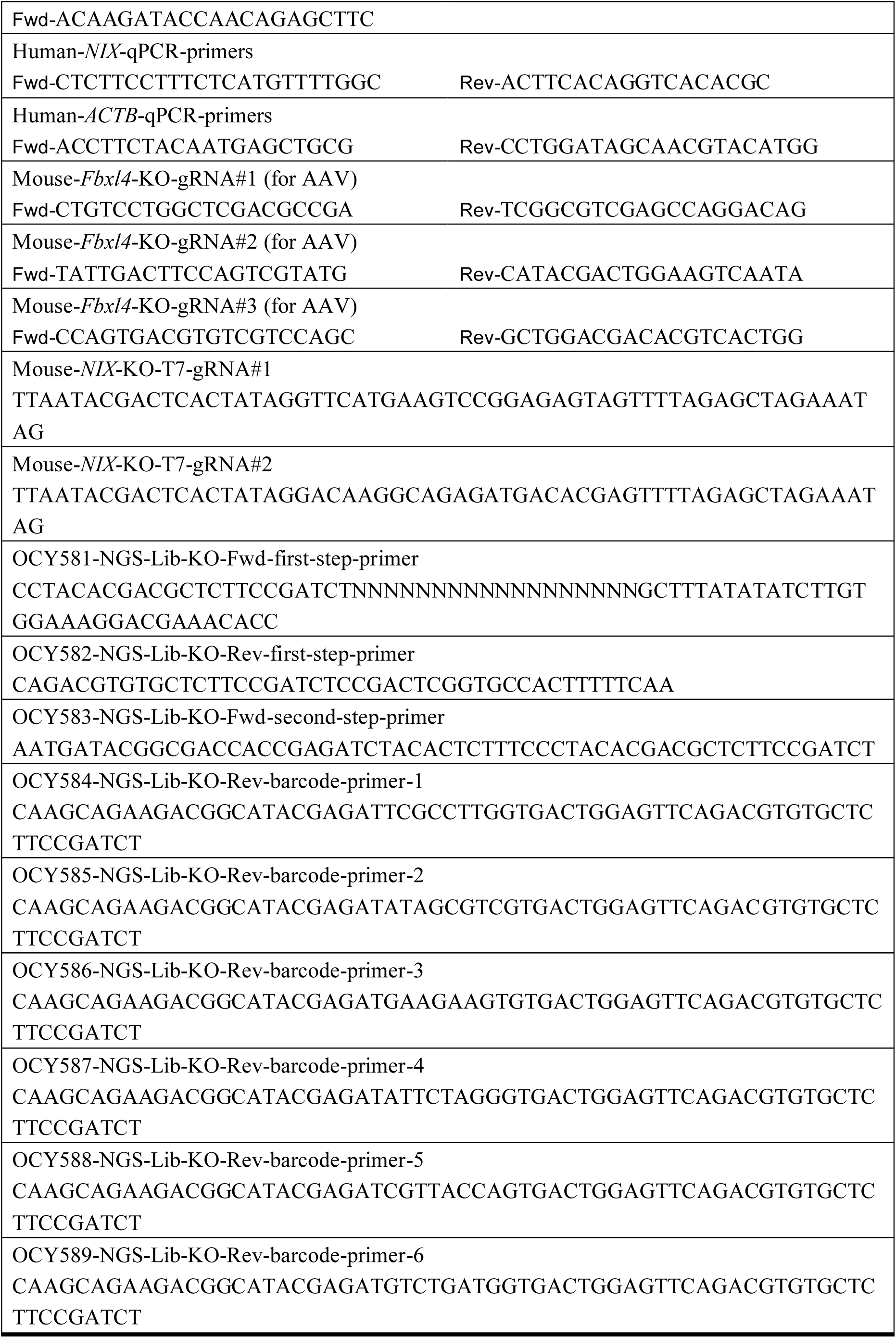
Oligonucleotides used in this study. Related to Methods.

## REFERENCES

Allen GFG, Toth R, James J, Ganley IG (2013) Loss of iron triggers PINK1/Parkin-independent mitophagy. EMBO reports 14: 1127–1135

Alsina D, Lytovchenko O, Schab A, Atanassov I, Schober FA, Jiang M, Koolmeister C, Wedell A, Taylor RW, Wredenberg A, Larsson N-G (2020) FBXL4 deficiency increases mitochondrial removal by autophagy. EMBO Molecular Medicine 12: e11659

Andreux PA, Blanco-Bose W, Ryu D, Burdet F, Ibberson M, Aebischer P, Auwerx J, Singh A, Rinsch C (2019) The mitophagy activator urolithin A is safe and induces a molecular signature of improved mitochondrial and cellular health in humans. Nature Metabolism 1: 595–603

Bai C, Sen P, Hofmann K, Ma L, Goebl M, Harper JW, Elledge SJ (1996) SKP1 Connects Cell Cycle Regulators to the Ubiquitin Proteolysis Machinery through a Novel Motif, the F-Box. Cell 86: 263–274

Barøy T, Pedurupillay CRJ, Bliksrud YT, Rasmussen M, Holmgren A, Vigeland MD, Hughes T, Brink M, Rodenburg R, Nedregaard B, Strømme P, Frengen E, Misceo D (2016) A novel mutation in FBXL4 in a Norwegian child with encephalomyopathic mitochondrial DNA depletion syndrome 13. European Journal of Medical Genetics 59: 342–346

Bellot G, Garcia-Medina R, Gounon P, Chiche J, Roux D, Pouysségur J, Mazure Nathalie M (2009) Hypoxia-Induced Autophagy Is Mediated through Hypoxia-Inducible Factor Induction of BNIP3 and BNIP3L via Their BH3 Domains. Molecular and Cellular Biology 29: 2570–2581

Bonnen Penelope E, Yarham John W, Besse A, Wu P, Faqeih Eissa A, Al-Asmari Ali M, Saleh Mohammad AM, Eyaid W, Hadeel A, He L, Smith F, Yau S, Simcox Eve M, Miwa S, Donti T, Abu-Amero Khaled K, Wong L-J, Craigen William J, Graham Brett H, Scott Kenneth L et al. (2013) Mutations in FBXL4 Cause Mitochondrial Encephalopathy and a Disorder of Mitochondrial DNA Maintenance. The American Journal of Human Genetics 93: 471–481

El-Hattab AW, Dai H, Almannai M, Wang J, Faqeih EA, Al Asmari A, Saleh MAM, Elamin MAO, Alfadhel M, Alkuraya FS, Hashem M, Aldosary MS, Almass R, Almutairi FB, Alsagob M, Al-Owain M, Al-Sharfa S, Al-Hassnan ZN, Rahbeeni Z, Al-Muhaizea MA et al. (2017) Molecular and clinical spectra of FBXL4 deficiency. Human Mutation 38: 1649–1659

Finck BN, Kelly DP (2006) PGC-1 coactivators: inducible regulators of energy metabolism in health and disease. The Journal of Clinical Investigation 116: 615–622

Fujiki Y, Hubbard AL, Fowler S, Lazarow PB (1982) Isolation of intracellular membranes by means of sodium carbonate treatment: application to endoplasmic reticulum. Journal of Cell Biology 93: 97–102

Gai X, Ghezzi D, Johnson Mark A, Biagosch Caroline A, Shamseldin Hanan E, Haack Tobias B, Reyes A, Tsukikawa M, Sheldon Claire A, Srinivasan S, Gorza M, Kremer Laura S, Wieland T, Strom Tim M, Polyak E, Place E, Consugar M, Ostrovsky J, Vidoni S, Robinson Alan J et al. (2013) Mutations in FBXL4, Encoding a Mitochondrial Protein, Cause Early-Onset Mitochondrial Encephalomyopathy. The American Journal of Human Genetics 93: 482–495

Gross A (2016) BCL-2 family proteins as regulators of mitochondria metabolism. Biochimica et Biophysica Acta (BBA) - Bioenergetics 1857: 1243–1246

Guna A, Stevens TA, Inglis AJ, Replogle JM, Esantsi TK, Muthukumar G, Shaffer KCL, Wang ML, Pogson AN, Jones JJ, Lomenick B, Chou T-F, Weissman JS, Voorhees RM (2022) MTCH2 is a mitochondrial outer membrane protein insertase. Science 378: 317–322

He Y-L, Gong S-H, Cheng X, Zhao M, Zhao T, Zhao Y-Q, Fan M, Zhu L-L, Wu L-Y (2020) BNIP3 phosphorylation by JNK1/2 promotes mitophagy via enhancing its stability under hypoxia. bioRxiv: 2020.08.27.271270

Hertz Nicholas T, Berthet A, Sos Martin L, Thorn Kurt S, Burlingame Al L, Nakamura K, Shokat Kevan M (2013) A Neo-Substrate that Amplifies Catalytic Activity of Parkinson’s-Disease-Related Kinase PINK1. Cell 154: 737–747

Huemer M, Karall D, Schossig A, Abdenur JE, Al Jasmi F, Biagosch C, Distelmaier F, Freisinger P, Graham BH, Haack TB, Hauser N, Hertecant J, Ebrahimi-Fakhari D, Konstantopoulou V, Leydiker K, Lourenco CM, Scholl-Bürgi S, Wilichowski E, Wolf NI, Wortmann SB et al. (2015) Clinical, morphological, biochemical, imaging and outcome parameters in 21 individuals with mitochondrial maintenance defect related to FBXL4 mutations. Journal of Inherited Metabolic Disease 38: 905–914

Jin J, Cardozo T, Lovering RC, Elledge SJ, Pagano M, Harper JW (2004) Systematic analysis and nomenclature of mammalian F-box proteins. Genes & Development 18: 2573–2580

Jumper J, Evans R, Pritzel A, Green T, Figurnov M, Ronneberger O, Tunyasuvunakool K, Bates R, Žídek A, Potapenko A, Bridgland A, Meyer C, Kohl SAA, Ballard AJ, Cowie A, Romera-Paredes B, Nikolov S, Jain R, Adler J, Back T, et al. (2021) Highly accurate protein structure prediction with AlphaFold. Nature 596: 583–589

Katayama H, Kogure T, Mizushima N, Yoshimori T, Miyawaki A (2011) A Sensitive and Quantitative Technique for Detecting Autophagic Events Based on Lysosomal Delivery. Chemistry & Biology 18: 1042–1052

Kim D-H, Sarbassov DD, Ali SM, King JE, Latek RR, Erdjument-Bromage H, Tempst P, Sabatini DM (2002) mTOR Interacts with Raptor to Form a Nutrient-Sensitive Complex that Signals to the Cell Growth Machinery. Cell 110: 163–175

Lavorato M, Nakamaru-Ogiso E, Mathew ND, Herman E, Shah N, Haroon S, Xiao R, Seiler C, Falk MJ (2022) Dichloroacetate improves mitochondrial function, physiology, and morphology in FBXL4 disease models. JCI Insight 7

Liang JR, Lingeman E, Ahmed S, Corn JE (2018) Atlastins remodel the endoplasmic reticulum for selective autophagy. Journal of Cell Biology 217: 3354–3367

Liu L, Feng D, Chen G, Chen M, Zheng Q, Song P, Ma Q, Zhu C, Wang R, Qi W, Huang L, Xue P, Li B, Wang X, Jin H, Wang J, Yang F, Liu P, Zhu Y, Sui S et al. (2012) Mitochondrial outer-membrane protein FUNDC1 mediates hypoxia-induced mitophagy in mammalian cells. Nature Cell Biology 14: 177–185

Liu L, Li Y, Wang J, Zhang D, Wu H, Li W, Wei H, Ta N, Fan Y, Liu Y, Wang X, Wang J, Pan X, Liao X, Zhu Y, Chen Q (2021a) Mitophagy receptor FUNDC1 is regulated by PGC-1α/NRF1 to fine tune mitochondrial homeostasis. EMBO reports 22: e50629

Liu S, Fu S, Wang G, Cao Y, Li L, Li X, Yang J, Li N, Shan Y, Cao Y, Ma Y, Dong M, Liu Q, Jiang H (2021b) Glycerol-3-phosphate biosynthesis regenerates cytosolic NAD+ to alleviate mitochondrial disease. Cell Metabolism 33: 1974–1987.e9

Liu Y-T, Sliter DA, Shammas MK, Huang X, Wang C, Calvelli H, Maric DS, Narendra DP (2021c) Mt-Keima detects PINK1-PRKN mitophagy in vivo with greater sensitivity than mito-QC. Autophagy 17: 3753–3762

Mammucari C, Milan G, Romanello V, Masiero E, Rudolf R, Del Piccolo P, Burden SJ, Di Lisi R, Sandri C, Zhao J, Goldberg AL, Schiaffino S, Sandri M (2007) FoxO3 Controls Autophagy in Skeletal Muscle In Vivo. Cell Metabolism 6: 458–471

McWilliams TG, Prescott AR, Allen GFG, Tamjar J, Munson MJ, Thomson C, Muqit MMK, Ganley IG (2016) mito-QC illuminates mitophagy and mitochondrial architecture in vivo. Journal of Cell Biology 214: 333–345

McWilliams TG, Prescott AR, Montava-Garriga L, Ball G, Singh F, Barini E, Muqit MMK, Brooks SP, Ganley IG (2018) Basal Mitophagy Occurs Independently of PINK1 in Mouse Tissues of High Metabolic Demand. Cell Metabolism 27: 439–449.e5

Mizushima N, Komatsu M (2011) Autophagy: Renovation of Cells and Tissues. Cell 147: 728–741

Moehlman AT, Youle RJ (2020) Mitochondrial Quality Control and Restraining Innate Immunity. Annual Review of Cell and Developmental Biology 36: 265–289

Narendra D, Tanaka A, Suen D-F, Youle RJ (2008) Parkin is recruited selectively to impaired mitochondria and promotes their autophagy. Journal of Cell Biology 183: 795–803

Narendra DP, Jin SM, Tanaka A, Suen D-F, Gautier CA, Shen J, Cookson MR, Youle RJ (2010) PINK1 Is Selectively Stabilized on Impaired Mitochondria to Activate Parkin. PLOS Biology 8: e1000298

Onishi M, Yamano K, Sato M, Matsuda N, Okamoto K (2021) Molecular mechanisms and physiological functions of mitophagy. The EMBO Journal 40: e104705

Palikaras K, Lionaki E, Tavernarakis N (2015) Coordination of mitophagy and mitochondrial biogenesis during ageing in C. elegans. Nature 521: 525–528

Petroski MD, Deshaies RJ (2005) Function and regulation of cullin–RING ubiquitin ligases. Nature Reviews Molecular Cell Biology 6: 9–20

Pickrell Alicia M, Youle Richard J (2015) The Roles of PINK1, Parkin, and Mitochondrial Fidelity in Parkinson’s Disease. Neuron 85: 257–273

Platt Randall J, Chen S, Zhou Y, Yim Michael J, Swiech L, Kempton Hannah R, Dahlman James E, Parnas O, Eisenhaure Thomas M, Jovanovic M, Graham Daniel B, Jhunjhunwala S, Heidenreich M, Xavier Ramnik J, Langer R, Anderson Daniel G, Hacohen N, Regev A, Feng G, Sharp Phillip A et al. (2014) CRISPR-Cas9 Knockin Mice for Genome Editing and Cancer Modeling. Cell 159: 440–455

Rapaport D (2003) Finding the right organelle. EMBO reports 4: 948–952

Ryu D, Mouchiroud L, Andreux PA, Katsyuba E, Moullan N, Nicolet-dit-Félix AA, Williams EG, Jha P, Lo Sasso G, Huzard D, Aebischer P, Sandi C, Rinsch C, Auwerx J (2016) Urolithin A induces mitophagy and prolongs lifespan in C. elegans and increases muscle function in rodents. Nature Medicine 22: 879–888

Sandoval H, Thiagarajan P, Dasgupta SK, Schumacher A, Prchal JT, Chen M, Wang J (2008) Essential role for Nix in autophagic maturation of erythroid cells. Nature 454: 232–235

Schweers RL, Zhang J, Randall MS, Loyd MR, Li W, Dorsey FC, Kundu M, Opferman JT, Cleveland JL, Miller JL, Ney PA (2007) NIX is required for programmed mitochondrial clearance during reticulocyte maturation. Proceedings of the National Academy of Sciences 104: 19500–19505

Spiegelman BM (2007) Transcriptional Control of Mitochondrial Energy Metabolism through the PGC1 Coactivators. In Mitochondrial Biology: New Perspectives, pp 60–69.

Sun N, Yun J, Liu J, Malide D, Liu C, Rovira Ilsa I, Holmström Kira M, Fergusson Maria M, Yoo Young H, Combs Christian A, Finkel T (2015) Measuring In Vivo Mitophagy. Molecular Cell 60: 685–696

Vara-Pérez M, Rossi M, Van den Haute C, Maes H, Sassano ML, Venkataramani V, Michalke B, Romano E, Rillaerts K, Garg AD, Schepkens C, Bosisio FM, Wouters J, Oliveira AI, Vangheluwe P, Annaert W, Swinnen JV, Colet JM, van den Oord JJ, Fendt S-M, et al. (2021) BNIP3 promotes HIF-1α-driven melanoma growth by curbing intracellular iron homeostasis. The EMBO Journal 40: e106214

Wu K, Fuchs Serge Y, Chen A, Tan P, Gomez C, Ronai Ze, Pan Z-Q (2000) The SCFHOS/β-TRCP-ROC1 E3 Ubiquitin Ligase Utilizes Two Distinct Domains within CUL1 for Substrate Targeting and Ubiquitin Ligation. Molecular and Cellular Biology 20: 1382–1393

Yao X, Wang X, Hu X, Liu Z, Liu J, Zhou H, Shen X, Wei Y, Huang Z, Ying W, Wang Y, Nie Y-H, Zhang C-C, Li S, Cheng L, Wang Q, Wu Y, Huang P, Sun Q, Shi L et al. (2017) Homology-mediated end joining-based targeted integration using CRISPR/Cas9. Cell Research 27: 801–814

Youle RJ, Narendra DP (2011) Mechanisms of mitophagy. Nature Reviews Molecular Cell Biology 12: 9–14

Zhang H, Bosch-Marce M, Shimoda LA, Tan YS, Baek JH, Wesley JB, Gonzalez FJ, Semenza GL (2008) Mitochondrial Autophagy Is an HIF-1-dependent Adaptive Metabolic Response to Hypoxia*. Journal of Biological Chemistry 283: 10892–10903

Zhang L-N, Zhou H-Y, Fu Y-Y, Li Y-Y, Wu F, Gu M, Wu L-Y, Xia C-M, Dong T-C, Li J-Y, Shen J-K, Li J (2013) Novel Small-Molecule PGC-1α Transcriptional Regulator With Beneficial Effects on Diabetic db/db Mice. Diabetes 62: 1297–1307

Zheng J, Cao Y, Yang J, Jiang H (2022) UBXD8 mediates mitochondria-associated degradation to restrain apoptosis and mitophagy. EMBO reports 23: e54859

